# Behavior emerges from unstructured muscle activity in response to neuromodulation

**DOI:** 10.1101/2021.03.10.434785

**Authors:** Amicia D. Elliott, Adama Berndt, Matthew Houpert, Snehashis Roy, Robert L. Scott, Carson C. Chow, Hari Shroff, Benjamin H. White

## Abstract

Identifying neural substrates of behavior requires defining actions in terms that map onto brain activity. Brain and muscle activity naturally correlate via the output of motor neurons, but apart from simple movements it has been difficult to define behavior in terms of muscle contractions. By mapping the musculature of the pupal fruit fly and comprehensively imaging muscle activation at single cell resolution, we here describe a multiphasic behavioral sequence in *Drosophila*. Our characterization identifies a previously undescribed behavioral phase and permits extraction of major movements by a convolutional neural network. We deconstruct movements into a syllabary of co-active muscles and identify specific syllables that are sensitive to neuromodulatory manipulations. We find that muscle activity shows considerable variability, which reduces upon neuromodulation. Our work provides a platform for studying whole-animal behavior, quantifying its variability across multiple spatiotemporal scales, and analyzing its neuromodulatory regulation at cellular resolution.

## Introduction

A major goal of neuroscience is explaining how nervous systems generate and organize behavior. This requires describing behavior in terms that can be correlated with neural activity. The dynamics of brain activity can be observed in whole brains at single cell resolution (Ahrens et al., 2013; Ardiel et al., 2017; Cong et al., 2017; Lemon et al., 2015; Nguyen et al., 2016; Pulver et al., 2015), but behavioral dynamics has not been captured at a similar level of detail (Datta et al., 2019). Progress in fine-mapping natural behavior, or “computational ethology” (Anderson and Perona, 2014), has benefited from recent advances in visual tracking (Johnson et al., 2020), 3D imaging (Hong et al., 2015), machine vision (Dankert et al., 2009), machine learning (Kabra et al., 2013; Machado et al., 2015), and image feature extraction (Berman et al., 2014; Mathis et al., 2018; Wiltschko et al., 2015). The primary focus has been kinematic, seeking to define anatomical movements at higher resolution. However, movements are complex products of motor neuron activity that balance agonist and antagonist muscle contractions and also provide anatomical rigidity. Here, we bridge the gap by detailing muscle activity at the single cell level.

Comprehensively monitoring muscle activity in behaving animals is achievable with genetically-encoded Ca^++^ indicators and has been demonstrated at single cell resolution in hydra (Szymanski and Yuste, 2019), roundworms (Ardiel et al., 2017), and larval fruitflies (Heckscher et al., 2012; Zarin et al., 2019). However, the application of muscle Ca^++^ imaging to characterize more complex sequences has been constrained by the challenge of tracking behavior in freely moving animals. This problem is resolved in pupal fruit flies where behavior is restricted to the puparium (Kim et al., 2006), which can be clarified for optical access to the confined animal. The pupa maintains a fixed position and orientation during movement, and all behavior is executed within a delimited field of view. Pupal Ca^++^ activity imaging has been demonstrated using pan-muscularly expressed GCaMP6s by Diao et al. (2017), who showed that the bulk Ca^++^ signal collected over ventral muscles exhibits temporal patterns that conform well to known body wall movements.

The behavioral hallmark of pupal development is the well-studied ecdysis sequence, which facilitates the transformation of the body plan at metamorphosis (Diao et al., 2017; Kim et al., 2015; Kim et al., 2006; Lahr et al., 2012; Mena et al., 2016; Zitnan and Adams, 2012). It is initiated after a long period of behavioral quiescence by the peripherally-released Ecdysis Triggering Hormone (ETH). Among the targets of ETH are neuroendocrine cells that express other neuromodulatory factors, including the hormones CCAP and Bursicon. Cells expressing Bursicon and CCAP become active approximately 10 min after the onset of pupal ecdysis and release of these hormones mediates the transition to the next behavioral phase. How nervous systems transform hormonal signals into temporally ordered motor sequences is an open question.

We use body-wide fluorescence imaging from the dorsal, lateral, and ventral views to characterize the pupal ecdysis sequence at single cell resolution. Using improved imaging methods—including a new pan-muscle LexA driver that permits dual imaging of muscle and neuron activity—we identify novel elements of the pupal ecdysis sequence, including undescribed movements and a phase of stochastic muscle activity preceding ecdysis. We find that muscle activity exhibits a high degree of variability, with individual muscles recruited stochastically into repeating small ensembles, which we call syllables. Syllable activity is synchronized over anatomical compartments to form movements, which are sufficiently stereotyped to be learned by a convolutional neural network (CNN). We can prevent synchronization at specific motor program transitions by suppressing neuromodulatory neurons, which is lethal. The suppression of proprioceptive neurons, which blocks initiation of pupal ecdysis, is also unexpectedly lethal. Overall, our analysis at single-cell resolution reveals a dynamical system in which movements are not rigidly specified but form from variable components subject to neuromodulatory reorganization.

## Results

### Pupal behavior described at cellular resolution

Although Drosophila behavior has been recorded at single-cell resolution, the drivers used to make such recordings express weakly at the pupal stage and cannot be used with Gal4 drivers targeting cells such as neurons. We identified a striated-muscle-specific gene, *l(2)01289* (aka CG9432), that expresses robustly from early larval stages through adulthood, which we call *hulk (hlk)*. We generated a Trojan exon insertion in the *hlk* gene that co-expresses LexA:QF and combined it with the Ca^++^ biosensor LexAop-GCaMP6s to report muscle activity. With this *hlk*-LexA>LexA_op_-GCaMP6s line (i.e. *hlk*>GCaMP6s), we imaged muscle Ca^++^ activity through the clarified puparium for approximately 90 minutes to capture pupal ecdysis behavior, *in vivo* (Methods, Fig. S1A, B).

Pupal ecdysis has three principal phases (Diao et al., 2017; Kim et al., 2006, Fig. 1A). The first phase (P1) consists of sustained longitudinal compression (“lifting”) of posterior abdominal segments accompanied by anteriorly-directed “rolling” contractions of the dorsal body wall that alternate left-to-right. The second (P2) features left-to-right alternating lateral “swinging” movements formed by unilateral, anteriorly-directed contractions, while the third (P3) consists of alternating left-right posteriorly-directed contractions that change into bilaterally symmetric contractions. These phases always proceed in the same order. The movements increase internal pressure to evert the head (i.e. force it out of the body cavity) and push the developing legs and wings to the body surface and elongate them (Zdarek and Friedman, 1986). Phalloidin labeling demonstrates that approximately half of larval muscles persist until pupal ecdysis, and all retain innervation by Ib synapses (Prokop, 2006, Fig. 1B-E). The most prominent loss of muscles occurs in the ventral and posterior compartments. Only 5 of 13 larval ventral muscles survive (Fig. 1D vs 1E), and one of these (M12) is absent in posterior segments (Table S1). Also missing from posterior segments are muscles M4 and M5. All five dorsal longitudinal muscles, except M4, are present in all segments, as are five of the six lateral transverse muscles. This was consistent across 17 animals, indicating that pupal ecdysis is executed by a standard set of persistent larval muscles.

**Figure 1.**
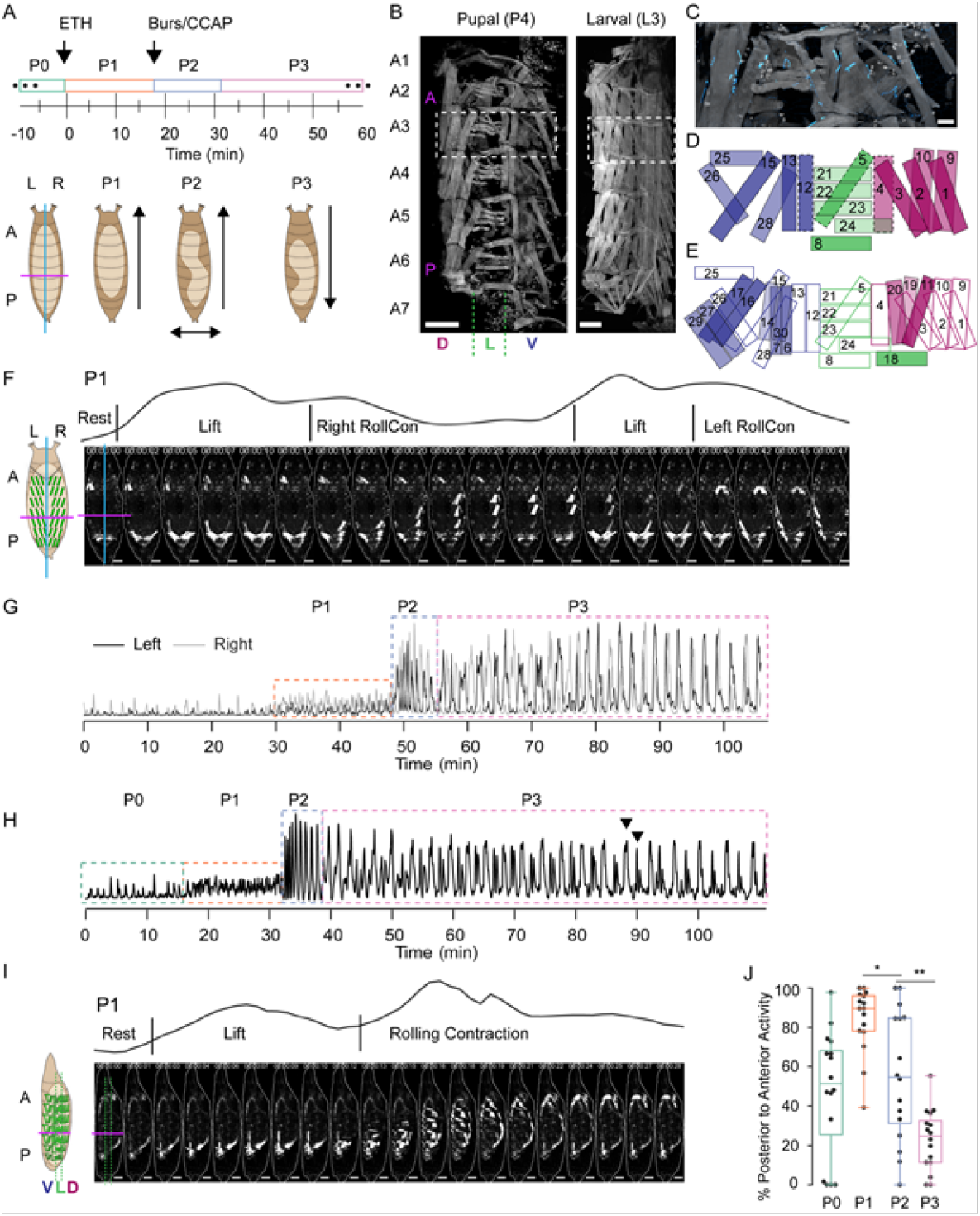
Pupal Ecdysis Behavior at Cellular Resolution. **(A)** Timeline of pupal ecdysis with phases indicated (top). Schematics show puparium (brown) and pupa (beige) from dorsal view (Left-Right axis, cyan; Anterior-Posterior boundary, magenta). Characteristic pupal movements for each phase indicated on right. Arrows, direction of movements. **(B)** Pupal and larval musculature stained with phalloidin (gray) showing one hemisegment (HS), midline to the right. HS A1-A7 are labeled (A3, boxed), as are axial compartments: A, anterior; P, posterior; D, dorsal; V, ventral; L, lateral. Scale bar, 250 μm. **(C)** Pupal muscles of HS A3 stained with phalloidin (gray) and for postsynaptic marker DLG1 (cyan). Scale bar, 25 μm. **(D)** Pupal muscles in a representative HS (e.g. dotted box in (B), left; based on 17 fillets). Muscles are labeled and coded blue (ventral), green (lateral), fuchsia (dorsal). Dotted lines indicate those present only in anterior segments. **(E)** Larval muscles in a representative HS (e.g. dotted box in (B), right; based on 6 fillets). Filled muscles degrade before pupal ecdysis. Colors as in D. **(F)** Muscle activity in a P1 bout (dorsal view) from a pupa expressing *hlk*>GCaMP6s. Schematic on left shows L-R and A-P axes as in (A). Cyan, dorsal midline; Magenta, A-P boundary between segments A4 and A5. Trace above images shows bulk Ca^++^ activity signal with movements (see Table S2). Scale bar, 250 μm. **(G)** Time traces of bulk Ca^++^ activity on the left (black) and right (gray) sides of a pupa executing the pupal ecdysis sequence and imaged from the dorsal side, as in (F). Alternating activity is evident. Dotted boxes, ecdysis phases. **(H)** Time trace of bulk Ca^++^ activity from a pupa imaged from the lateral side. Dotted boxes, ecdysis phases. Arrowheads, peak-double peak bouts of late P3. **(I)** Muscle activity in a P1 bout (lateral view) from a pupa expressing *hlk*>GCaMP6s. Schematic on left shows A-P partition between HSA4-A5 (magenta) and lateral muscle compartments (dotted green). Trace above images shows bulk Ca^++^ activity signal with movements (see Table S2). Scale bar, 250 μm. **(J)** Bouts in each phase (%) with P-to-A activity, such as shown in I. N=16 pupae. * p<0.05; **, p<0.01. **See also Figure S1, Table S1, and Movies S1 and S2**.

The muscle activity patterns of animals imaged from the dorsal side match known behaviors, such as the bilateral posterior “lifts” and left-right alternating “rolling contractions” of P1 (Fig. 1F, Movie S1). Temporal patterns of bulk Ca^++^ activity differentiated phases P1, P2, and P3 (Fig. 1G). Traces show phase-specific oscillations of varying amplitude and frequency, with individual oscillations conforming to bouts of movement (see Methods). Consistent with (Diao et al., 2017), the alternating left-right oscillations of P1 persisted through P2 and P3, with coordinated bilateral activity becoming dominant only in P3. Bulk Ca^++^ activity imaged from the lateral side showed the same three phases (Fig. 1H), and oscillations reflected bouts that included identifiable movements (Fig. 1I). The lift is performed by nearly all muscles across the dorsal-ventral (D-V) axis in the posterior segments, while rolling contractions (RollCons) lack activity in the ventral longitudinal muscles 12, 13, and 15 (Movie S2). Posterior-to-anterior (P-to-A) waves of activity in P1 reverse direction after head eversion in P2 (Fig. 1J), as previously shown (Kim et al., 2006). These data confirm and refine previous observations and demonstrate that the *hlk*>GCaMP6s line accurately reports the ecdysis sequence behaviors.

### Muscle imaging reveals an initial phase of random neurogenic activity

Ca^++^ imaging revealed a phase of random muscle activity prior to the onset of pupal ecdysis (Fig. 1H, Movie S3), which we call Phase 0 (P0). It begins approximately 3 hours before P1 and divides into distinct bouts of muscle activation (Fig. 2A, Movie S3). Individual muscle length changes during P0 are small (≤ 25%) compared to P1-P3 (25-40%; Fig. S2A). To determine if P0 muscle activation is myogenic or neurogenic, we created a dual-reporter fly line with *hlk-LexA* driving expression of the red fluorescent Ca^++^ biosensor jRGECO in the muscle and VGlut-Gal4 driving a Synaptotagmin-GCaMP6s fusion protein (Syt-GCaMP6s) in motor neurons, where it localizes to the neuromuscular junction (NMJ, Fig. 2B). *In vivo* imaging revealed synaptic Ca^++^ activity at the NMJ 30-40 minutes prior to the first muscle Ca^++^ response (Fig. 2C). As P0 progresses, coincident synaptic and muscle activity increases until the onset of P1, when nearly all muscles and their synaptic inputs are synchronously active (Fig. 2C, D) except M12, which remains unresponsive to input until P2 (Fig. S2B). Near complete muscle responsiveness may serve as a checkpoint for starting the ecdysis sequence and is possibly implemented by proprioceptive feedback. Class I dendritic arbor (da) neurons, dmd1, vbd, and dbd act as proprioceptors during larval locomotion (Vaadia et al., 2019) and remain present at the pupal stage (Fig. 2E). Moreover, bulk Ca^++^ activity in sensory neurons is correlated with muscle activity during P0 (Fig. 2F), and sensory neurons commonly show correlated Ca^++^ activity with adjacent muscles (Fig. 2G). When class I da neurons were suppressed with UAS-Kir2.1, Ca^++^ became sustained and widely distributed during P0 before twitching stopped (Fig. S2C, D). Unexpectedly, all animals died before P1 (n=10).

**Figure 2.**
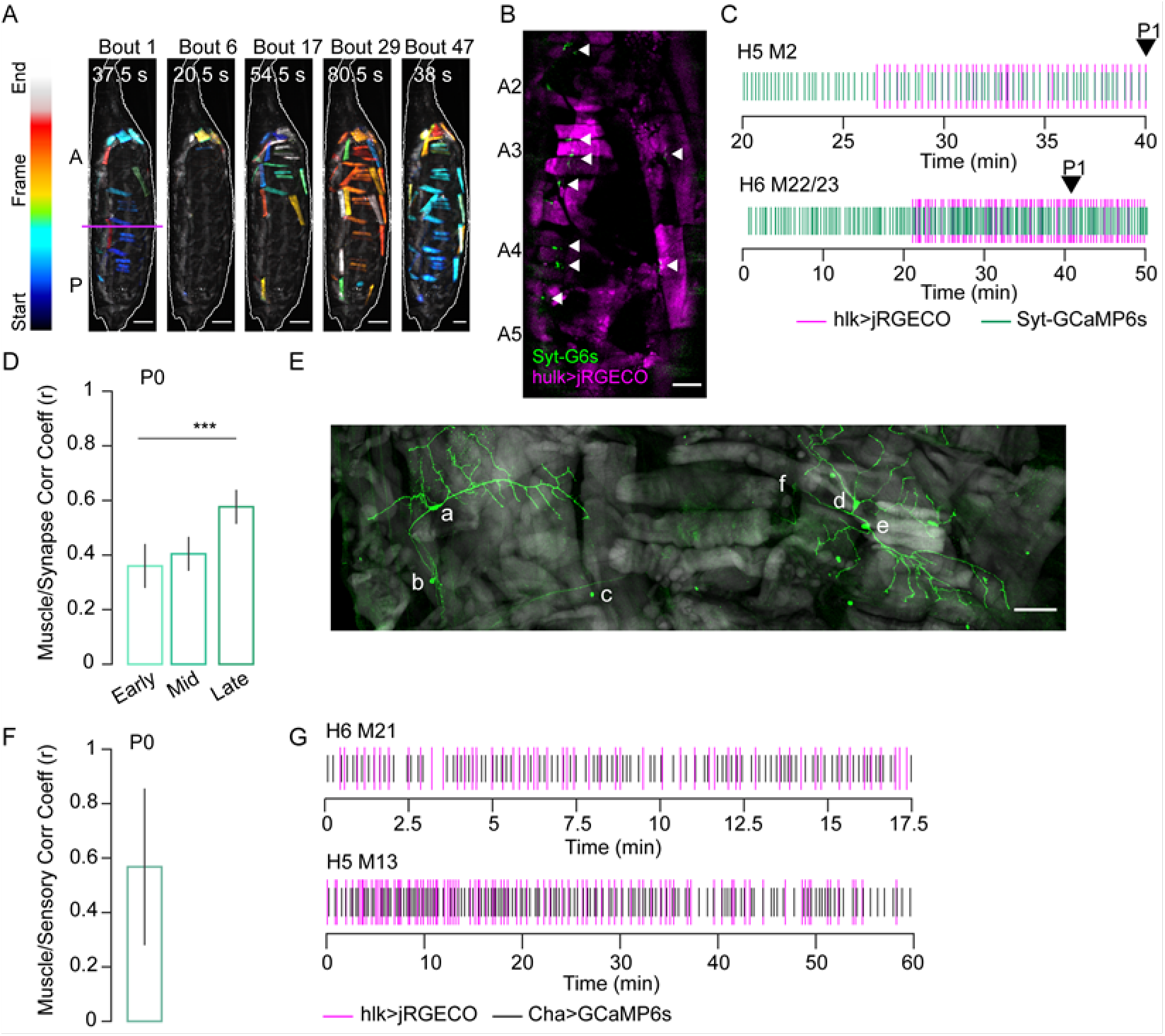
Stochastic Muscle Activity Precedes the Ecdysis Sequence. **(A)** Time-coded projections of muscle Ca^++^ activity (lateral view) in 5 P0 bouts. Muscle activity is distinct for each bout. Bout durations as indicated; image frames were color-coded according to color scale (left). Magenta line, A-P boundary. Scale bar, 250 μm. **(B)** Muscle (magenta; *hlk*>jRGECO) and neuromuscular junction (NMJ, green; *VGlut*>Syt-GCaMP6s) activity in body wall HS A2-A5. Arrows, active NMJs. Scale bar, 200 μm. **(C)** Representative raster plots generated from peaks in *hlk*>jRGECO (magenta) and *VGlut*>Syt-GCaMP6s (green) activity for the indicated muscles and their respective NMJs. Arrow, P1 onset. **(D)** Pearson correlation coefficients for *VGlut*>Syt-GCaMP6sand *hlk*>jRGECO activity peaks in multiple muscle/NMJ pairs during early, mid, and late temporal bins, relative to P1 onset. Early P0, N=19 muscles; mid P0, N=89; late P0, N=35. ***, p≤0.001. **(E)** HS A3 from pupa expressing mCD8-GFP (green) in class I da neurons. Phalloidin-stained muscles (gray). Neuronal somata: a: vpda, b: vbd, c: dbd, d: ddaD neuron, e: ddaE, f: dmd1. Scale bar, 50 μm. **(F)** Pearson correlation coefficient for bulk Ca^++^ activity peaks in muscles labeled with *hlk*>jRGECO and sensory neurons labeled with ChaT-GCaMP6s. N=10 pupae. **(G)** Rasters compare Ca^++^ activity peaks of the indicated muscle (magenta; *hlk*>jRGECO) with those in an adjacent sensory neuron (black; *ChaT*-GCaMP6s) during P0. See also Figure S2 and Movie S3.

### Muscle activity patterns identify elementary movements

The transition from P0 to P1 is accompanied by the first overt movements, with bouts typically consisting of a Lift followed by a RollCon. The transition from P1 to P2 is demarcated by a sudden behavioral switch in which a P1 bout is followed by four to five bouts containing only a Swing. We used changing muscle activity patterns with associated body wall displacements to define five further canonical movements, all executed in characteristic anatomical compartments (Fig. 3A, Table S2). We also define a precise onset for P3, which had previously been difficult (see for example Diao et al., 2017; Kim et al., 2006).

**Figure 3.**
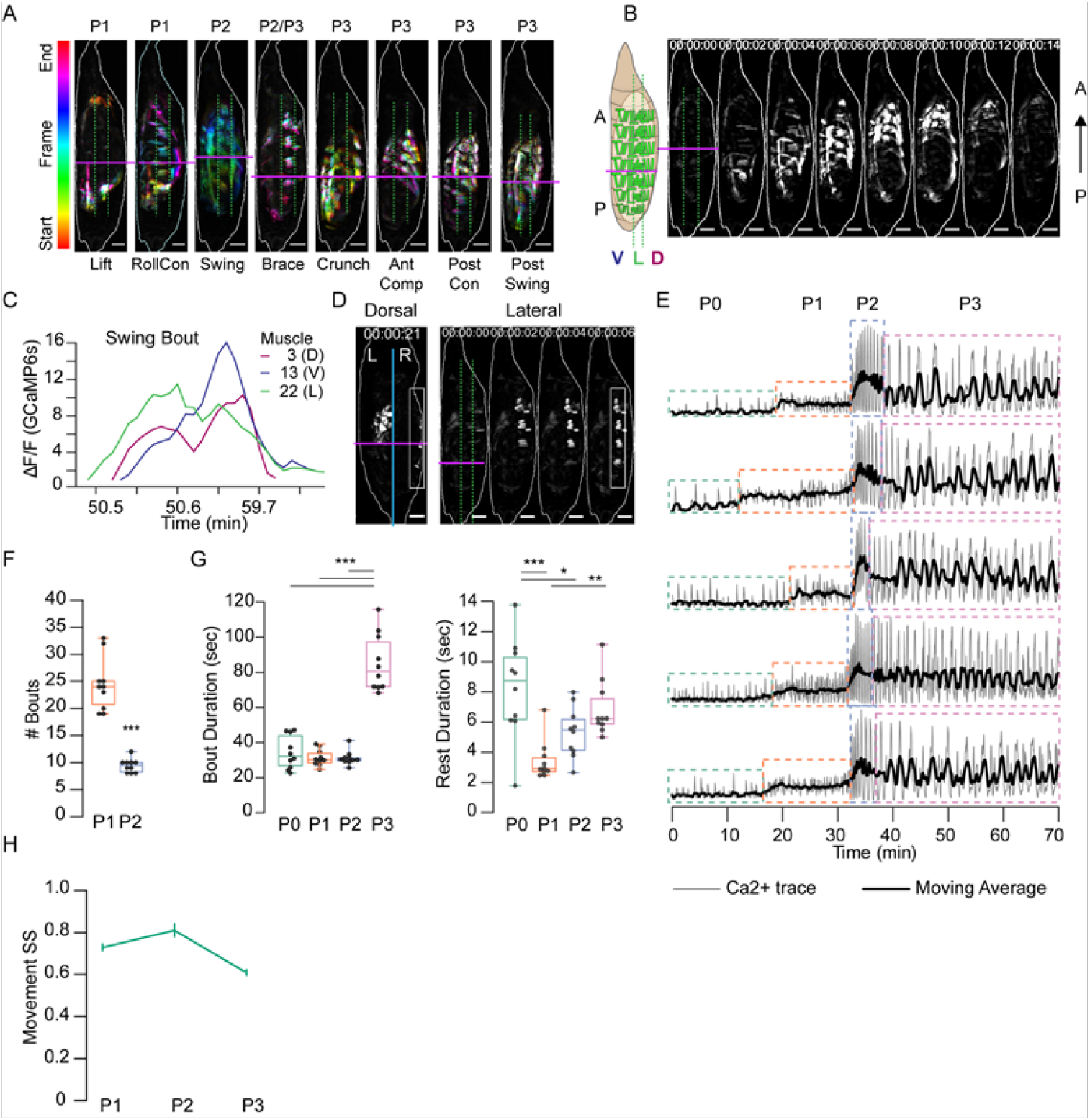
Muscle Activity Patterns Identify Elementary Movements. **(A)** Time-coded projections of muscle activity (lateral view) executed during labeled movements of the indicated phases. Magenta line, A-P boundary; dotted green lines, lateral compartment. Scale bars, 250 μm. **(B)** Muscle activity comprising a Swing movement showing the P-to-Awave of activation across dorsal, lateral, and ventral compartments. Schematic and first image on left indicate the anterior (A) posterior (P) boundary (magenta line), and the lateral compartment (green dotted lines). Scale bars, 250 μm. **(C)** Representative Ca^++^ traces from a single Swing bout measured for M3, M13, and M22 in HSA4 show coincident activity across the dorsal (D), lateral (L), and ventral (V) compartments. **(D)** Dorsal and lateral P2 muscle Ca^++^ activity during the Brace. White boxes, active lateral Brace muscles, coincident with swing activity on the opposite side of the animal (dorsal view). Cyan, dorsal midline. Scale bars, 250 μm. **(E)** Muscle Ca^++^ traces for 5 representative pupae imaged from the lateral side. Gray, bulk activity; black, 100 frame moving average. Phases as indicated. **(F)** Number of manually annotated activity bouts in P1 and P2. N=10 pupae. ***, p≤0.001. **(G)** Bout (left) and interbout-interval (right, “Rest”) durations, for manually annotated pupae in (F). *, p<0.05; **, p≤0.01; ***, p≤0.001. (H) SequenceMatcher similarity scores (SS) for movement sequences of Pl, P2, and P3. See also Figure S3,Tables S1 and S2, and Movie S4.

Muscle activity in a Swing is coordinated across the dorsoventral compartments as it travels anteriorly (Fig. 3B, C). While initial Swings are rapid, they slow after head eversion (Kim et al., 2006) and are then accompanied by a movement we call the “Brace.” (Fig. 3A, D). The Brace is performed by concurrent contraction of lateral transverse muscles M21-23 and M8 in anterior hemisegments, followed by contraction of these same muscles in posterior hemisegments (Fig. 3D, lateral view). The Brace begins the shift of activity from P-to-A waves in P1 to A-to-P waves in P3. Onset of P3 is indicated by a movement we call the “Crunch” (Fig. 3A). The first Crunch follows the last P2 bout after a relatively long interbout interval and combines ventral contractions in the posterior compartment with dorsal contractions in the anterior compartment. The compartmentalized and complex movements that follow the Crunch include what we call the “Anterior Compression” (AntComp), “Posterior Contraction” (PostCon), and “Posterior Swing” (PostSwing). The first two comprise what have been termed “stretch compressions” (Kim et al., 2006). All of these movements are unique to P3.

Each of the elementary movements defined in Fig. 3A is associated with the activity of specific muscles, and we trained a CNN to recognize and annotate them (Fig. S3A, B; Movie S4). Using the CNN to measure movement durations (Fig. S3C), together with measurements of bout and phase durations (see Methods), we characterized the variability of pupal ecdysis behavior at the level of phases, bouts, and movements.

### Behavioral stereotypy increases with spatiotemporal level of description

The relative stereotypy of the pupal ecdysis sequence can be seen from bulk Ca^++^ traces (Fig. 3E) but variation exists in the duration and bout number of P1 and P2 (Table S3, Fig. 3F), and in the bout durations of all phases (Fig. 3G). Coefficients of variation (CV) were lowest for P2 for all phase and bout parameters examined (Table S3). This finding is consistent with the developmental importance of P2 and suggests that its execution is the most tightly regulated of all the phases. Movement durations also showed variability, with CVs exceeding 50% (Table S3). However, the order in which movements were executed as determined by the SequenceMatcher algorithm (see Methods) indicated considerable stereotypy. Sequence similarity scores (SS) for movements were computed pairwise for all bouts within each animal by phase and compared across animals. The mean SSs of the sequences for P1, P2, and P3 were all above 0.6, a threshold for similarity (Fig. 3H), with CVs of 20-44%. P3 had the lowest SS and P2 the highest.

To evaluate the stereotypy of the muscle activation patterns used to generate individual movements, we also used the SequenceMatcher algorithm. The order in which muscles were activated in bouts of P1 yielded an SS of 0.44 ± 0.17, indicating low similarity. However, this SS differed significantly from that of shuffled sequences (0.32 ± 0.12, p<0.001). The sequences are thus not entirely random. Variability was evident between animals (Fig. 4A) and for individual P1 movements across animals (Fig. 4B). The low similarity of muscle activity patterns suggested that movements may be generated by muscles that are consistently active together even when their individual activation times vary between bouts. Co-active muscle groups have been shown to be important for larval locomotion (Zarin et al., 2019); we thus identified groups of muscles for which: a) all muscles were co-active in three or more consecutive frames, b) the group was identified in at least 80% of the movements in a given animal and, c) the group was identified in at least 80% of animals. We found eight co-active muscle groups, which we call “pupal muscle ensembles” (PMEs; Fig. 4D). Muscles forming a PME are not recruited in a consistent order (Fig. 4C). In addition, we found several muscles that individually satisfied criteria b) and c), but not as part of a group. Collectively with the PMEs, we call these movement-associated muscles “syllables,” and those active in the 8 pupal ecdysis movements are listed in Table S2.

**Figure 4.**
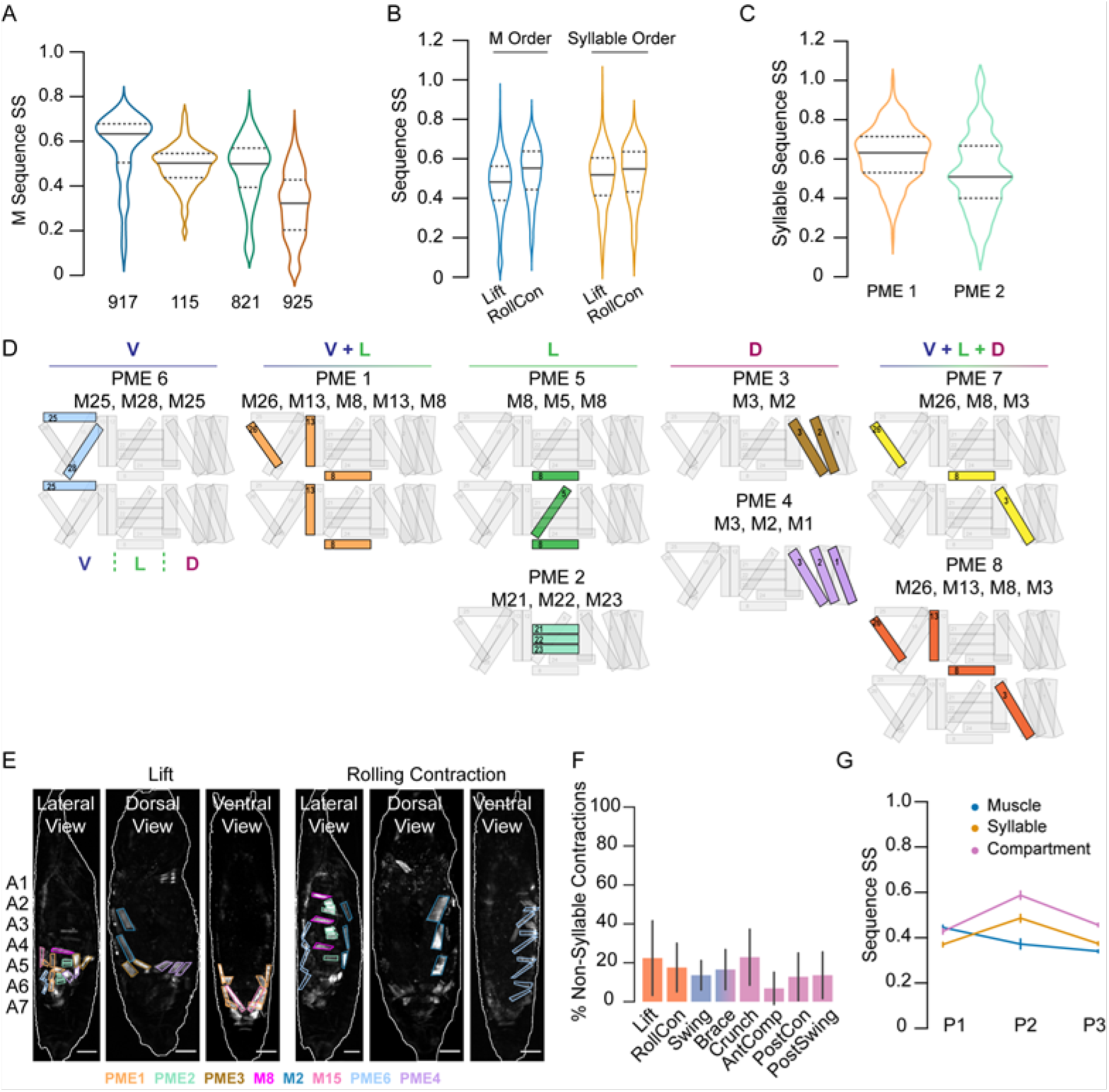
Coactive Muscles Define Movement Syllables. **(A-C)** istribution of similarity scores (SS) for single-muscle (M) activation sequences or syllable sequences in 4 manually annotated *hlk*>GCaMP6s pupae. **(A)** M activation sequences in P1 bouts by pupa (numbers). **(B)** Single-muscle activation sequences (blue plots) and syllable sequences (orange plots) in P1 movements across pupae. **(C)** Single-muscle activation sequences in PME1 and PME2. **(D)** Pupal muscle ensembles shown schematically on hemisegment musculature and organized by anatomical compartment; active muscles colored. **(E)** Syll ables associated with Lift and RollCon muscle activity: lateral, dorsal, and ventral views. Scale bars, 250 μm. **(F)** Percentage of muscle activations not annotated as part of a syllable, color-coded by phase. N=4 animals. **(G)** Mean SS (± SD) for activation sequences of compartments (pink), syllables (orange), and single muscles (blue) for each phase by bout. N=4 pupae. See also Table S2.

Surprisingly, the patterns of activation of the 5 syllables associated with P1 movements (Fig. 4E) showed SSs similar to those obtained using the muscle sequences (Fig. 4B, orange plots). This suggests that random muscle activations outside of syllables may contribute to the P1 movements, and such idiosyncratic activations were common in movements of all phases (Fig. 4F). However, unlike the syllable sequences of P1 movements, those of P2 and P3 movements were significantly more consistent than the sequence of recruited individual muscles (Fig. 4G). Because syllables are often confined to particular anatomical compartments, we also compared the activity sequences in the D-V and A-P compartments across P1-P3. P1 bouts exhibit only modest similarity but the bouts of P2 and P3 are intermediate in similarity to those of movements and syllables (compare Figs. 4G and 3H). Thus, the observed stereotypy of motor execution in pupal ecdysis is highest for phases, and incrementally decreases with spatiotemporal scale. The least stereotypy is seen in the rank order of recruitment of individual muscles, which is less consistent than the activation of syllables.

### Muscle mechanics provide insight into movement composition and purpose

To quantify how Ca^++^ activity in muscles of the pupal syllabary generates movement, we measured the normalized maximum shortening (ΔL/L) and peak fluorescence intensity (ΔF/F) for each muscle contraction in segments A3-A5. Increases in fluorescence moderately correlated with muscle shortening (r=-0.54; Fig. S4A). To determine which muscles shorten the body wall, we calculated the average ΔL/L for each muscle over each phase, focusing first on P2 because of its role in promoting morphological change. For P2, M12 and the muscles comprising PMEs 1 (M26, M13, M8), 2 (M21-23), 3 (M2, M3), and 4 (M1-M3) shorten the most (Fig. 5A). In each hemisegment, these contractions compress the animal longitudinally and along the D-V axis, with a wave of such compressions traversing the body wall in the P-to-A direction during the Swing (Fig. 5C; Movie S4). The greatest constriction across the D-V axis occurs in posterior segments, consistent with pronounced shortening in PME2 muscles (M21-23) in A5. Progressively decreased shortening of PME2 muscles is observed in A4 and A3. Shortening of the ventral and dorsal longitudinal muscles (M12, M13, M1-3) is more uniform across hemisegments, but the absence of M12 in posterior hemisegments and somewhat greater shortening of the dorsal longitudinal muscles anteriorly is consistent with the greater longitudinal compression of anterior hemisegments (Fig 5C).

**Figure 5.**
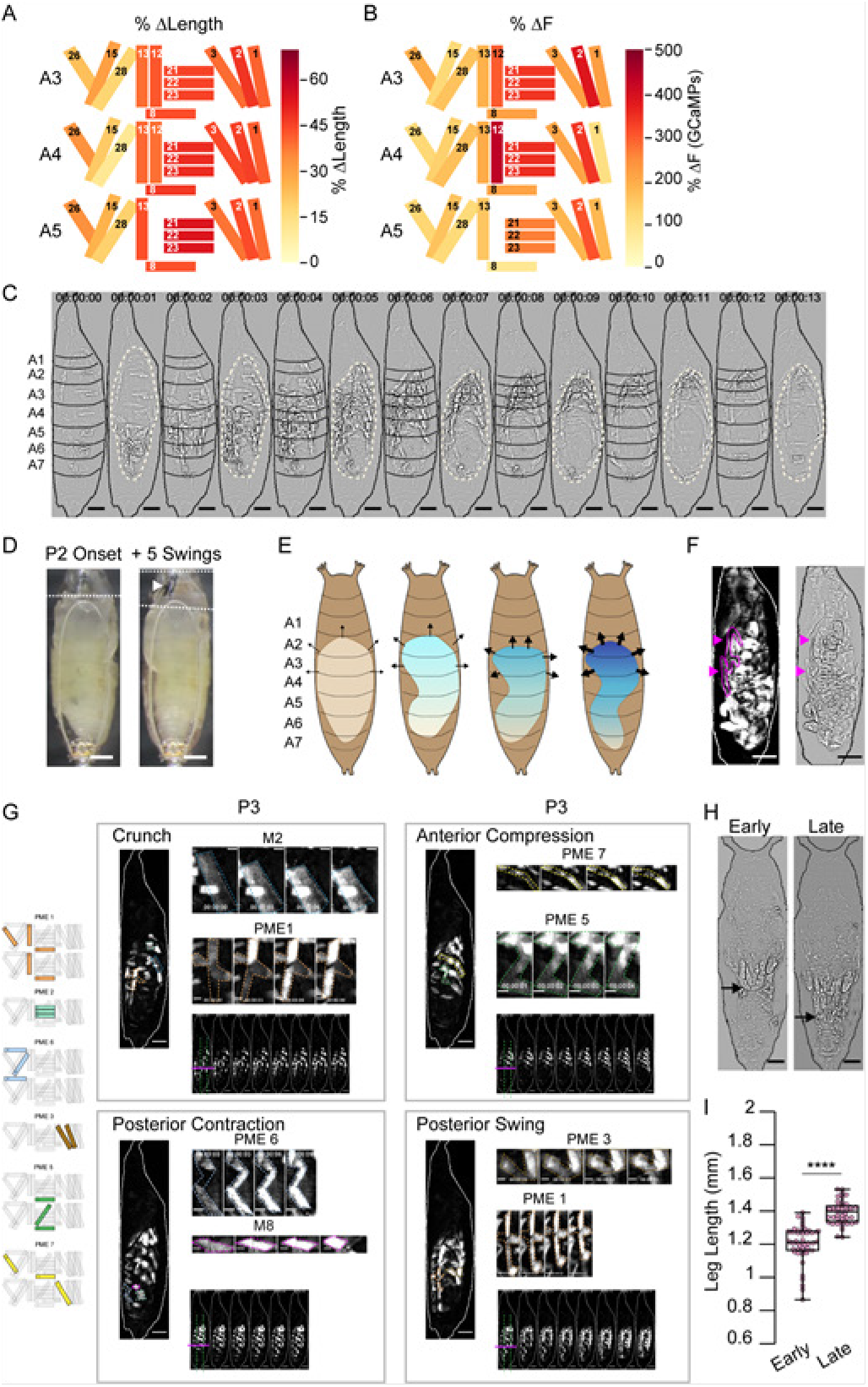
Muscle Mechanics during P2 and P3 Behaviors. **(A-B)** Changes in muscle properties during contraction of indicated muscles in HS A3-A5 during P2. **(A)** Change in muscle fiber length (ΔL/L). **(B)** Change in GCaMP6s fluorescence (ΔF/F). % changes were calculated from values at activity onset and at maximum activity for N=24 pupae and color-coded as indicated. **(C)** Muscle activity of a Swing after Laplace transform to show body wall distortion during movement (black, HS boundaries; beige outlines, pupal body). Scale bars, 250 μm. **(D)** Images before (left) and after (right) head eversion. Dotted lines indicate body anterior prior to and after head eversion. Head, black arrow; larval mouth hooks, white arrow. Scale bars, 200 μm. **(E)** Illustrated effects of the Swing movement on internal pressure. The P-to-A bending of the body wall pushes hemolymph forward, increasing pressure (blue gradient and arrows) in the anterior compartment. Darker blue and thicker arrows, increased pressure. **(F)** Larval dorsal tracheal trunks (magenta arrows) are deposited on the puparium wall during P2 Swings by a pupa expressing tdTomato in muscles. Trachea are visualized by scattered light from fluorescent muscles (grayscale, left) and highlighted in magenta in Laplace transformed image (right) for clarity. Scale bar, 250 μm. **(G)** Muscle activity underlying P3 movements. Each box contains: a time-series montage of the activity (bottom; lateral compartment and A-P boundary indicated); a single-frame image (left) with principal syllables outlined (see key, far left); and montages showing the temporal evolution of syllables. Scale bars, 250 μm (pupal images); 50 μm (montages). **(H)** Leg extension during P3 imaged from the ventral side. Ends of the extending legs (arrows) are shown at early and late P3. **(I)** Leg length during the first (Early) and last (Late) 10 bouts recorded of P3. Scale bars, 250 μm. ***, p≤0.0001. See also Figure S4, Table S3, Movie S4.

Comparing ΔL/L with the corresponding average (ΔF/F) reveals hemisegmental differences (Fig. 5B vs. Fig. 5A) including an increase in peak Ca^++^ activity of PME2 muscles (M21-23), moving from A5 to A3 (Fig. 5B). This trend runs opposite to muscle shortening, which means that in successive anterior hemisegments, the PME2 muscles work harder to generate a smaller length change. This suggests counterforces on the anterior body wall, consistent with increased pressure to evert the head (Fig. 5D) as has been measured in blowflies (Zdarek and Friedman, 1986). Each Swing is initiated posteriorly when the hemolymph is uniformly distributed throughout the body cavity and internal pressure is low. Ascending compression of the body wall on one side pushes the opposite side of the body against the static puparium. This prevents further body wall distension on that side and the compression wave drives hemolymph forward, like squeezing a tube of toothpaste from the bottom up. This creates pressure anteriorly which pushes the head out (Fig. 5E). While a bilaterally coordinated compression might evert the head more efficiently, the unilateral Swing has a second function: It extrudes the larval tracheal linings on each side of the body. These are deposited in ascending segments on the puparium with each Swing (Fig. 5F).

The two movements of P1 share features of the Swing and likely prepare the animal for P2. The Lifts draw out the dorsal tracheal trunks, which remain attached to the posterior spiracles (Robertson, 1936); the RollCons may help fragment the linings of the stretched trunks in anterior segments so that they are efficiently extruded at P2. Essential to the Lift is compaction of the posterior hemisegments, which is accomplished by bilateral contraction of almost the same syllables as the Swing (Table S2). The RollCon, like the Swing, is performed unilaterally in a P-to-A direction. However, it fails to compact hemisegments as it traverses them and only deflects the dorsal body wall. Single-muscle changes in ΔL/L and ΔF/F underlying RollCons are much smaller than those for the Swing (compare Fig. S4B, C with Fig. 5A, B). In addition, RollCons engage only a subset of the syllables comprising the Swing (Table S2). Rare Ca^++^ activity of M12 is idiosyncratic in P1 and does not contribute to ΔL/L. The coordinated contractions of M12 with other muscles during the Swing, together with the recruitment of additional syllables and higher Ca^++^ activities, explain why RollCons result only in deflections of the dorsal body wall while Swings cause hemisegmental compaction to bend the entire animal (compare Fig. S4D with Fig. 5C). Overall, the movements of P1 appear to merge in P2, combining into one robust concerted P-to-A movement. In this view, the Swing is a unilateral Lift that successfully propagates into the anterior compartment.

In P3, coordinated movement across anatomical compartments separates into multiple compartmentalized movements, which form activity patterns without strictly repeating units. Bulk Ca^++^ activity imaged from the lateral side shows an initial oscillatory pattern of variable frequency and amplitude that evolves into a more fixed pattern of alternating wide peaks and slim double peaks (see Fig. 1I, arrows). The initial variable period of activity is heralded by the Crunch (Fig. 5G), which is generated by the contralateral activation of PME1 in ventral hemisegments posterior to segment 4 and M2 in anterior dorsal segments. These contractions slightly lift the posterior segments and compact the anterior segments. The Crunch is typically followed by a Brace or an AntComp (Fig. 5G). The latter movement compacts the dorsal and anterior compartments via contractions of PME7, PME3, and M1 and with ventral activity in PME6. Realignment of the body is achieved by the execution of a PostCon followed by a PostSwing (Fig. 5G), each composed of syllables in Table S2.

The block of movements containing a Crunch, Brace, AntComp, PostCon, and PostSwing yield an A-to-P flow of activity. They constitute a fairly regular repeating unit with some variation in the order. This block forms the double peaks seen in the bulk Ca^++^ trace and as P3 evolves, its variable repetition increasingly alternates with a modified block that lacks the PostSwing and is followed by long interbout intervals. In addition, later P3 blocks also include M12 contractions in PostSwings and sometimes Crunches, which visibly increase the compaction of the anterior segments. The increased compaction may drive hemolymph posteriorly into the appendages to lengthen them (Fig. 5H, I).

### Neuromodulators increase behavioral stereotypy

We used the inwardly rectifying K^+^ channel, Kir2.1, to selectively silence neurons that critically regulate entry into P1 and P2 (Diao et al., 2016; Kim et al., 2015). Specifically, we suppressed neurons expressing the B isoform of the ETH receptor (N_ETHRB_), and two overlapping populations of neurons expressing CCAP and Bursicon. The former manipulation disrupts P1 initiation by blocking abdominal lifting, while the latter blocks initiation of P2 (Diao et al., 2017). We monitored the effects on Ca^++^ activity using *hlk*>GCaMP6s.

In animals in which N_ETHRB_ neurons are suppressed, the baseline increase in bulk muscle Ca^++^ during P1 is severely attenuated relative to WT (Fig. 6A, B). Although PMEs 2, 3, and 6 characteristic of P1 (Table S2) appear 10-15 min prior to P2 (Fig. 6C), activity in M15 and M1 (and thus PME4) is missing. Muscles of PME1 are also not simultaneously active and thus do not exhibit ensemble activity (Fig. 6C, D). Finally, activity in the D-V compartments is not usually synchronized in the posterior segments. These data indicate that a principal population of ETH targeted neurons coordinates muscles into syllables to produce the lift movement. Components of the Lift also remain absent from later movements. For example, M1 activity remains disrupted during Swings (Fig. S5A).

**Figure 6.**
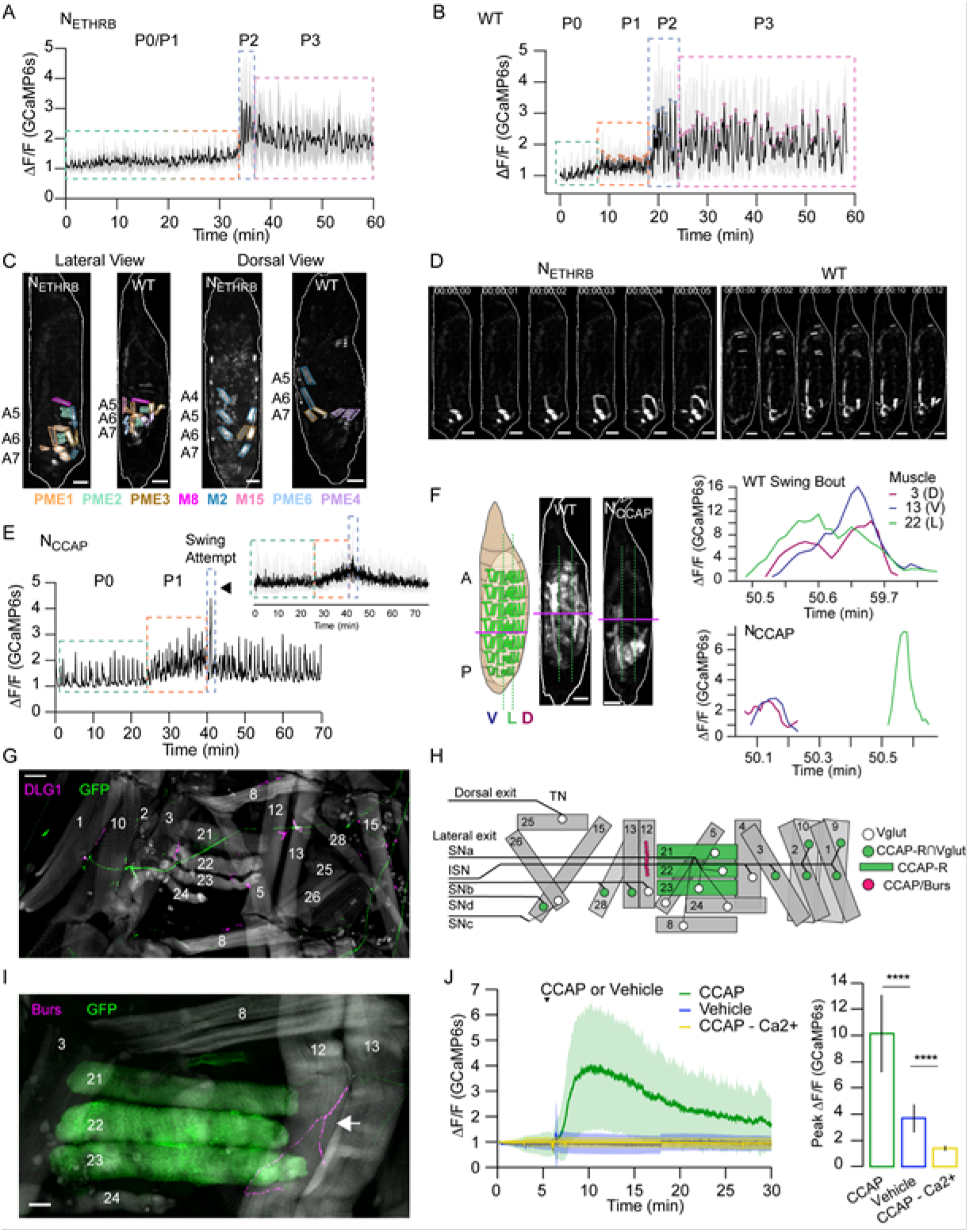
Neuromodulator Roles in Behavior. **(A-B)** Lateral view bulk Ca^++^ activity traces, mean (black), and SD (grey), for **(A)** animals with suppressed N_ETHRB_ (N=6), and **(B)** wild type pupae (N=16). Lack of change in activity precludes discrimination between PO and P1 for N_ETHRB_-suppressed animals. **(C)** Comparison of muscle activity in N_ETHRB_-suppressed (N_ETHRB_) and wildtype (WT) animals during the execution, or attempted execution, of P1 movements. Lateral view shows lift-like (N_ETHRB_) and Lift (WT) activity; dorsal view shows RollCons. Syllables are outlined. Scale bars, 250 μm. **(D)** Muscle activity in N_ETHRB_-suppressed and WT pupae shows disruption in posterior muscle activation. Scale bars, 250 μm. **(E)** Ca^++^ activity trace (lateral view) for animal with suppressed CCAP-expressing neurons (N_CCAP_). Inset, mean activity (± SD) for 10 N_CCAP_-suppressed pupae. Dotted boxes indicate identifiable phases. Blue box consists of a single partial swing-like movement where P2 typically begins (arrowhead). **(F)** Images of muscle activity during a Swing in WT or a partial swing-like movement in N_CCAP_-suppressed animals. Schematic on left indicates the A-P boundary (magenta) and lateral compartment (dotted green). On the right are Ca^++^ traces for M3, M13, and M22 in HS A4 from a single swing for a WT and N_CCAP_-suppressed animal. Co-incidence across the dorsal (D), lateral (L), and ventral (V) compartments is lost in N_CCAP_. Scale bars, 250 μm. **(G)** Motor axons expressing mCD8-GFP (green) under the control of the CCAP-R0 Vglut Split Gal4 driver innervate a subset of muscles (phalloidin, gray) at synapses stained for the postsynaptic marker DLG1 (magenta). HS A3 is shown. Scale bar, 50 μm. **(H)** Map of CCAP-R motor neuron innervation (green circles) of pupal muscles. Innervation by motor neurons expressing only VGlut, and not CCAP-R, (white circles) are also shown, as are muscles expressing CCAP-R (green rectangles) and the type III synapse on M12 that releases CCAP and Bursicon (fuchsia circles). (l)CCAP-R-expressing muscles M21-M23, visualized with CCAP-R-Gal4>mCD8-GFP (green) are located adjacent to the type III terminal on M12, which is immunopositive for CCAP and Bursicon (magenta, white arrow). HS A4 is shown. Gray, phalloidin-stained muscles. Scale bar, 50 μm. **(J)** CCAP application elicits robust Ca^++^ responses from M2l-M23 (green; mean ± SD) in live, filleted *hlk*>GCaMP6s animals. No response is seen with vehicle only (blue line; mean ± SD) or in Ca^++^-free media (yellow; mean ± SD). N=7-10 pupae, 12 HS/pupa. See also Figure S5,Table S1, and Movies S5 and S6.

Suppressing CCAP-secreting neurons (N_CCAP_) results in normal Ca^++^ activity during P0 and P1, but P2 and P3 activity is absent (Fig. 6E). Although repetitive P2 swinging is absent, a single, partial swing-like movement is observed after numerous P1 bouts, suggesting that the transition to P2 may be attempted but is not maintained (Movie S5). M1-3 activate asynchronously in anterior segments so that PME4 fails to form correctly. Consequently, the P-to-A wave on the dorsal side is disrupted by anterior contractions occurring too early (Fig. S5B). The partial swing propagates only through segment A5 and accompanying segmental compression is limited to A6 and A7 (Fig. S5C). The lateral muscles comprising PME2 are unsynchronized with the dorsal and ventral longitudinal muscles (Fig. 6F). Finally, the dorsal and ventral longitudinal muscles in segments A4-A5 change less in fluorescence (ΔF/F=157.2 ± 33.2) and length (ΔL=29.4 μm ± 6.23) than in WT (ΔF/F=272.3 ± 92.6; ΔL=42.2 μm ± 2.52). While N_CCAP_-suppression is lethal (Diao et al., 2016), there is persistent activity resembling P1 Lifts and RollCons with no significant reversal in P-to-A direction before death. There are no obvious transitions and our CNN detected few P3-specific movements (Fig. S5D). We conclude that N_CCAP_ modulates several aspects of the transition to P2, including: 1) generalized increase in muscle activity during P2; 2) coordination of syllables along the A-P axis that facilitates full body swings; and 3) coordination of activity across the D-V axis, as indicated by the desynchronization of PME2 activity.

The loss of synchronous activity across D-V compartments led us to investigate the innervation pattern of motor neurons that express the CCAP receptor (CCAP-R, Diao et al., 2017). Intersectional labeling of CCAP-R-expressing motor neurons using the Split Gal4 system (Luan et al., 2006) showed that these neurons innervate approximately half of the pupal muscles via Ib synapses, including all dorsal and three of the six ventral muscles (Fig. 6G, H). Although none of the motor neurons innervating the lateral transverse muscles express CCAP-R, the transverse muscles M21-23 of PME2 are labeled by the CCAP-R-Gal4 driver (Fig. 6H, I, Table S1). This suggests that CCAP centrally modulates motor neuron output to dorsal and ventral muscles, while directly modulating lateral transverse muscles. At the larval stage, CCAP is co-released with Bursicon from Type III terminals on muscles M12 and M13, which straddle the muscles of PME2 (Veverytsa and Allan, 2011). Anti-Bursicon staining established the persistence of Type III terminals on M12 (Fig. 6I, magenta). In addition, we confirmed the responsiveness of M21-23 to CCAP in fillet preparations treated with bath-applied peptide (Fig. 6J, Movie S6). Our results demonstrate a role for N_CCAP_ in coordinating syllable activity across both the A-P and D-V axes and a peripheral role for CCAP in directly modulating lateral muscles.

Genetic data suggest that CCAP and Bursicon act synergistically at pupal ecdysis (Lahr et al., 2012), and neurons expressing the Bursicon receptor have been identified as essential for ecdysis motor programs (Diao et al., 2017). Consistent with Bursicon’s colocalization with CCAP in central neurons, we find that suppressing the subset of Bursicon-expressing neurons (NBurs) has effects similar to N_CCAP_ suppression: P1 activity is normal, but P2 and P3 are not correctly executed (Fig. S5E) and the animals die without everting their heads. Animals also execute a single swing-like movement after numerous bouts of P1, but in this case it consists of an entire anteriorly-directed wave and some subsequent patterned activity is observed that resembles AntComp, Crunch, Brace, and PostSwing movements, which can be identified by our CNN (Fig. S5F). This activity lacks organization and remains desynchronized during the swing-like movement activity in the D-V compartments (Fig. S5G). Additional swing-like movements also occur, but always separated by other types of movement. We conclude that D-V synchronization for abdominal swinging requires NBurs, as does sustained execution of swinging behavior, and that full coordination of activity across the A-P axis additionally requires non-Bursicon-expressing neurons in N_CCAP_. The results of suppressing neuromodulatory signaling support both central and peripheral roles for the ecdysis hormones in promoting syllable coordination across the A-P and D-V axes. This coordination promotes the observed coherence of behavioral execution despite the variable timing of activation of individual muscles.

## Discussion

Behavior is linked to neural mechanisms by the muscle activity that governs movement. To gain insight into how nervous systems specify behavior, we examined muscle activity during the *Drosophila* pupal ecdysis sequence at single-cell resolution using genetic Ca^++^ indicators. The pupal ecdysis sequence consists of multiple motor programs, dependent for their execution on hormonal cues. We find that hormonal signaling coordinates muscle activity across individual muscle ensembles and anatomical compartments to ensure stereotypy of behavioral execution. Although stereotypy is evident at the level of phases, the recruitment of muscles into movements is not stereotyped and some muscle activity is not correlated with movement. Importantly, a phase of stochastic muscle Ca^++^ activity precedes the onset of behavior, indicating that prior to the action of ETH, nervous system activity exhibits intrinsic variability. This variability is reduced, but not eliminated, by the action of neuromodulators, which incrementally increase behavioral coherence.

### The emergence of stereotypy from variable muscle activity

Our results significantly extend previous descriptions of pupal ecdysis and illustrate the power of pan-muscle Ca^++^ imaging. Behavioral fine-mapping at single-cell resolution permits the definition and automated detection of elemental movements, the identification of a syllabary of movement-associated muscles and muscle ensembles, and the analysis of their sensitivity to neuronal manipulations (Fig. 7). Importantly, single-cell analysis permits the identification of muscle activity that is not consistently associated with movements. The most salient example of such idiosyncratic activity occurs in P0, a previously undescribed phase of muscle activity lacking coordinated movement. Variability persists in phases P1-P3, which exhibit idiosyncratic muscle activation comingled with stereotyped movement syllables. Furthermore, muscle recruitment into syllables, and recruitment of syllables into movements, exhibits considerable variability both within and across animals. All observations suggest that variability is a pervasive feature of the pupal ecdysis sequence with stereotypy emerging only at higher levels of description.

**Figure 7.**
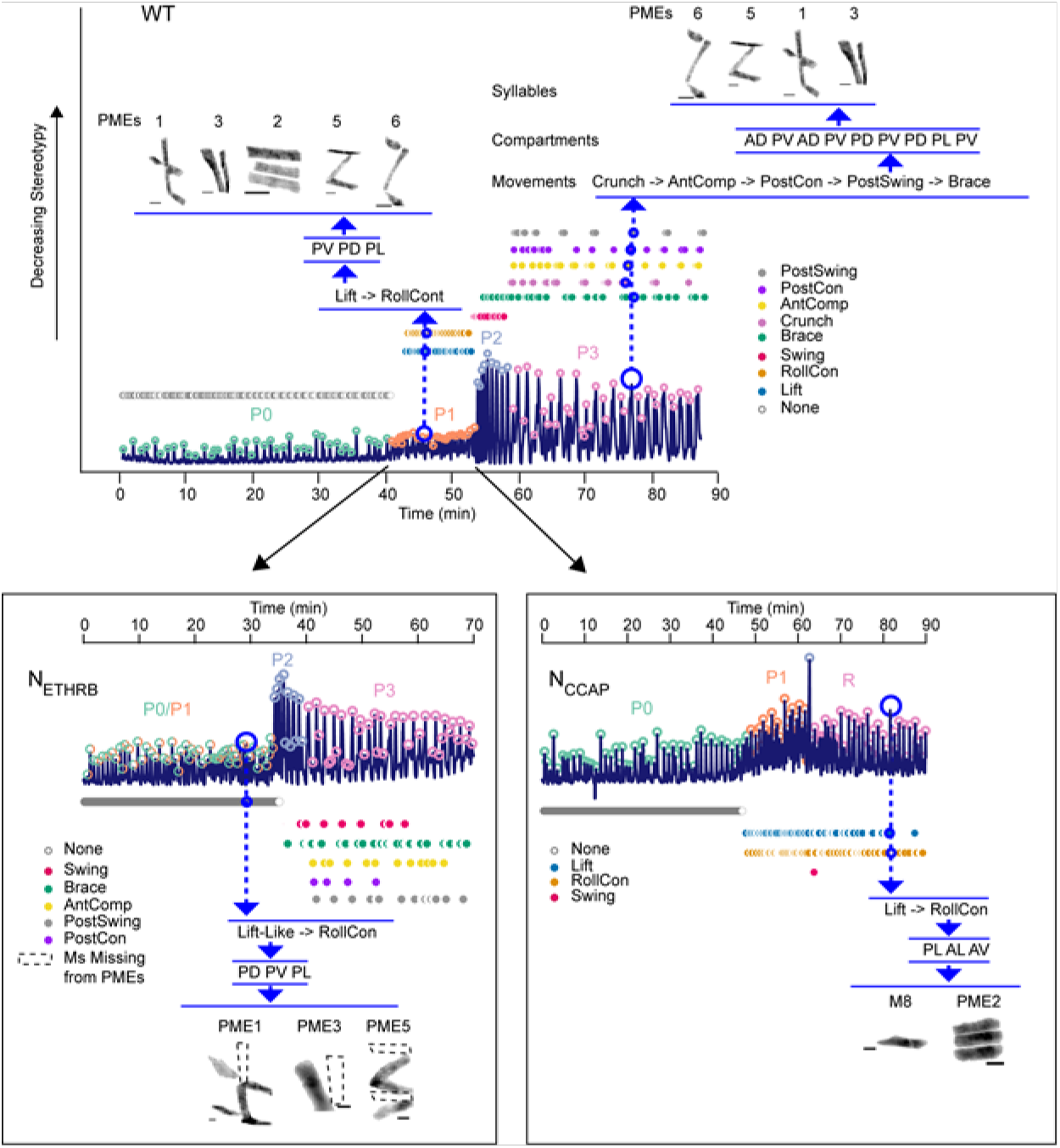
Decomposition of Pupal Ecdysis Behavior from Single Muscle Ca^++^ Activity. (Top) Representative bulk Ca^++^ activity over the four phases of the pupal ecdysis sequence, with behavioral bouts circled (P0, teal; Pl, orange; P2, blue; P3, pink). Identified movements (filled circles) are indicated. Upper right, movements during a P3 bout (large open blue circle) in order (small open blue circles) with the anatomical compartments of a PostSwing indicated in the order of initiation. PMEs recruited for the PostSwing are indicated above. Upper left, a P1 bout and Lift movement similarly represented. Bottom left, decomposed Ca^++^ trace for a N_ETHRB_-suppressed pupa. Dotted lines in PMEs indicate missing muscles. Bottom right, trace from N_CCAP_-suppressed pupa. Bout containing a Pl-like RollCon— executed where P3 would normally be observed— illustrated. Scale bars, 50 μm.

Because pupal ecdysis is independent of environmental factors and executed in the absence of competing physiological needs, it is likely that its variability is intrinsic to the ecdysis network. A possible source of variability is proprioceptive feedback, which in larvae has been shown to report body wall deformation (Vaadia et al., 2019) and regulate crawling speed (Hughes and Thomas, 2007)—roles consistent with sensory feedback providing timing cues to central pattern generators (Stein, 2014). We find that proprioceptive feedback is necessary for the initiation of the pupal ecdysis sequence, perhaps by reporting successful muscle activation in response to motor neuron input during P0. After this, feedback may not be essential since a fictive sequence is generated by an excised pupal brain in its absence (Diao et al., 2017; Kim et al., 2006; Mena et al., 2016). While proprioception may tune motor output during pupal ecdysis, the major source of variability seems to lie with the neural output pattern itself since stochasticity is already evident in P0 and then appears to extend to other phases.

### Neuromodulatory regulation of behavioral transitions and variability

Variability in pupal motor output is substantially altered by the neuromodulatory action of the ecdysis hormones ETH, CCAP, and Bursicon. Previous evidence indicates that these hormones induce the motor activity characteristic of P1 and P2, even in an isolated pupal nervous system (Diao et al., 2017; Kim et al., 2006). Consistent with the ability of neuromodulators to reconfigure motor networks (Marder, 2012), this induction likely represents reorganization and possibly stabilization of network activity, and also increased network coherence. P1 is distinguished from P0 by the presence of coherent movements and P2 is more coherent than P1 in recruitment of muscles, syllables, and compartmental activity. The reduced stereotypy in P3 may result from waning neuromodulatory action. The ecdysis hormones thus reorganize motor output and increase the stereotypy of its execution. This conclusion is supported by suppression of neuromodulatory signaling, which disrupts muscle activity at multiple levels, as expected from the broad distribution of ecdysis hormone receptors (Diao et al., 2017; Diao et al., 2016; Kim et al., 2006).

Expression of CCAP-R in both motor neurons and muscles suggests such broad-based, coordinating action. This receptor is expressed in motor neurons that innervate dorsal and ventral muscles. These motor neurons share synaptic inputs that are distinct from those innervating the lateral muscles (Landgraf et al., 2003; Zarin et al., 2019). However, lateral muscles M21-23, also express CCAP-R and likely receive CCAP from Type III synapses on M12. The pattern of CCAP-R expression may thus facilitate neuromuscular synchronization along the entire dorsoventral axis during the Swing movements of P2 when CCAP is released. Notably, swinging is not established when CCAP-expressing neurons are suppressed. Also, because M12 is present only in segments A1-A4, CCAP input to lateral muscles in the posterior compartment may be delayed. This may favor the late appearance of the Brace, which changes the character of the Swing after head eversion. The selective anterior retention of M12 and the ventral loss of much of the larval musculature illustrates how neuromodulation couples to anatomical asymmetries in compartmentalizing muscle activity across body axes. Neuromodulation may also explain the generalized failure of muscles to respond to synaptic input at the onset of P0 since *Drosophila* muscles are targets of a variety of neuropeptides including myoinhibitory peptides, which have been implicated in pupal ecdysis behavior (Kim et al., 2015).

What causes the release of Bursicon, CCAP, and ETH at pupal ecdysis is unknown, but our results suggest that ecdysis activity itself may be a factor. Release of ETH occurs only after muscle responsiveness to neural stimulation is ensured, suggesting that the latter may represent a checkpoint for release. Similarly, the aborted swing-like activity occurring in the absence of N_CCAP_ and NBurs activity indicates that entry into P2 has a hormone-independent component that may act as a checkpoint for hormone release, which then sustains P2 network activity. Checkpoint control mechanisms have been proposed to operate in the adult ecdysis sequence of locusts and crickets (Carlson, 1977; Hughes, 1980).

### Identifying neural determinants of behavior

A goal of computational neuroethology (Datta et al., 2019) is to describe behavior at a level of resolution that permits the identification of its neural determinants. The muscle-level description provided here lends itself naturally to this purpose. For example, the activity patterns of the muscles comprising PME2 have likely neural correlates within the synaptic and neuromodulatory networks that govern pupal ecdysis behavior. At the neuromodulatory level, the N_CCAP_ cells that terminate on M12 are likely sources of CCAP modulation of PME2 muscles, perhaps to help reverse their P-to-A activation at P2. At the synaptic level, the regular, intersegmental pattern of activation of PME2 muscles in nearly all pupal movements and behavioral bouts (Table S2) suggests that PME2 motor neurons are driven by CPG neurons that generate anteroposterior rhythms. Our results thus provide testable predictions about the patterns of neuromodulatory and synaptic connectivity between muscles, motor neurons, and premotor interneurons of various types.

Testing these predictions will be facilitated by data emerging from reconstruction of the larval CNS at synaptic resolution (Clark et al., 2018; Kohsaka et al., 2019; Zarin et al., 2019). Although pupal behaviors differ from those of the larva, broad similarities suggest that at least some neural substrates are shared. The initial P-to-A activity flow of the pupal ecdysis sequence is reminiscent of larval forward peristalsis and its reversal after alternating bilateral Swing movements resembles the switch to backward peristalsis following sensory stimuli (Carreira-Rosario et al., 2018; Tastekin et al., 2018). Importantly however, the basic features of larval locomotion are identifiable in the default activity of the excised larval brain (Lemon et al., 2015; Pulver et al., 2015), whereas the pupal nervous system produces patterned activity resembling pupal ecdysis only in response to ETH. The promise of pupal ecdysis as a behavioral model is in permitting investigation of the mechanisms by which intrinsic neuronal activity is organized by neuromodulatory control.

## Acknowledgments

We thank Dr. Feici Diao for advice and help initiating this project. We are grateful to the Bloomington Drosophila Stock Center (NIH P40OD018537) for fly lines used in this study and to Dr. Ted Usdin and the NIMH Systems Neuroscience Imaging Resource (ZIC-MH002963) for use of the Leica SP8 confocal microscope. This work was supported by the Intramural Research Programs of the NIGMS (FI2-GM117582, AE), NIMH (ZIA-MH002800, BW), NIDDK (CC), and the NIBIB (HS).

## Author Contributions

Conceptualization, A.D.E., H.S., C.C., and B.H.W; Methodology, A.D.E. and B.H.W.; Software, A. D.E., S.R., and A.B.; Formal Analysis, A.D.E., S.R., A.B., and M.H.; Investigation, A.D.E., A.B., and M.H.; Resources, R.L.S.; Writing – Original Draft, A.D.E. and B.H.W.; Writing – Review & Editing, A.D.E., H.S., C.C., A.B., M.H., R.L.S. and B.H.W; Funding Acquisition, A.D.E., H.S., C.C., B. H.W.; Visualization, A.D.E., B.H.W., H.S.; Supervision, A.D.E., H.S., C.C., and B.H.W.

## STAR Methods

### CONTACT FOR REAGENT AND RESOURCE SHARING

Further information and requests for resources and reagents should be directed to and will be fulfilled by the Lead Contact, Benjamin White.

### EXPERIMENTAL MODEL AND SUBJECT DETAILS

#### Drosophila stocks and rearing conditions

Vinegar flies of the species Drosophila melanogaster were used in this study. Flies were raised on cornmeal-molasses-yeast medium or Nutri-Fly German food (Genesee Scientific) and housed at 25°C and 65% humidity in a 12h light/dark cycle. Both males and females were used in this study and all experiments analyzed animals at stages L3 (third instar), P2 (Bainbridge and Bownes, 1981) ~6 hours after pupariation, or at the time of pupal ecdysis, ~12 h after pupariation. Fly stocks described in previous publications include: VGlut-Gal4DBD (i.e. VGlutMI04979-Gal4DBD) and ChaT-Gal4 from (Diao et al., 2015) and CCAPR-p65AD and CCAPR-Gal4 (Diao et al., 2017), ETHRB-Gal4 (Diao et al., 2016). *hlk* was identified by the larval muscle expression pattern of the NP3137-Gal4 enhancer trap line whose P{GawB} element is inserted at the 5’ end of the *l(2)01289* gene. To make the *hlk-T2A-LexA* line, PBS-KS-attb1-2-PT-SA-SD-1-T2A-LexA:QFAD-Hsp70 was injected into y w; l(2)01289^MI05738^/Cy; + embryos (Bloomington stock# 42105) and adults were screened for loss of the y+ marker. All injections were made by Rainbow Transgenic Flies, Inc (Camarillo, CA). The CCAP-Gal4 line was a kind gift from J. Ewer (Universidad de Valparaiso, Chile). All other fly lines, listed in the Key Resources Table, were obtained from the Bloomington Drosophila Stock Center at Indiana University.

### METHOD DETAILS

#### Immunohistochemistry

Third instar larvae and stage four pupae with an air bubble and visible gut movement indicating ecdysis proximity (Bainbridge and Bownes, 1981) were isolated from their vials, chilled for 20 minutes on ice, and placed into a dissection dish with 1X phosphate-buffered saline (1X PBS). Animals were pinned dorsal-side or ventral-side up (as indicated) at the anterior and posterior ends, a small incision was made along the entire dorsal or ventral midline, and the visceral organs were removed. Tissues were then fixed in 4% paraformaldehyde for 30 minutes at RT and washed in 1X PBS three times. A block was performed for 1 hour at RT in PBT (1X PBS, 0.5% Triton-X) with 5% Normal Goat Serum (NGS). The tissues were then incubated with Rabbit anti-GFP (Life Technologies, USA) diluted to 1:250 in PBT and mouse anti-DLG (Developmental Studies Hybridoma Bank, Iowa City, IA) at 1:500 for 48 hours at 4°C. After six 30-minute washes in PBT with shaking, samples were incubated with secondary antibodies Alexa Fluor 647, 488, and Alexa Fluor Phalloidin 555 (Life Technologies, USA) at 1:500 in PBT with 5% NGS for 48 hours at 4°C (Life Technologies, USA). Samples were washed again three times for 20-minute washes in 1X PBT with shaking and mounted on #1.5 glass coverslips with Prolong Diamond (Life Technologies, USA). Confocal imaging was performed using a Leica SP8 with AOBS, bidirectional resonant scanning, and a 20X/0.75 NA air objective. Unless otherwise noted, the images presented are maximum intensity projections, produced in Fiji ImageJ (Schindelin et al., 2012), of multi-point Z-stacks collected through the entire preparation.

#### Manipulation of Neuronal Activity

All neuronal suppression experiments were conducted using two copies of UAS-Kir2.1 in a parental line generated by combining insertions on Chromosomes II and III with *hlk*-LexA::QFAD and LexAOP-GCaMP6s, respectively.

#### Live fluorescence microscopy

##### Intact animals

P4 stage pupae were isolated as for immunohistochemistry, rinsed with 50% bleach for 3 min, rinsed in PBS, and immersed in a custom-chamber with 2,2-thiodiethanol (TDE, Sigma Aldrich), which rendered the puparium transparent and immobilized the animal with the dorsal, ventral, or lateral side facing the objective (Fig. S1A). Muscles only on the selected side were imaged using an epifluorescence stereomicroscope (Nikon SMZ25) with a 1x/0.3 NA objective, GFP filter cube, 4x magnification, low depth-of-field, and a partially closed aperture diaphragm. Fluorescence from muscles opposite the imaging field is effectively excluded because the pupal musculature lies superficially along the body wall and light is scattered by internal tissues. All muscle activity on one side could be rapidly captured in a single frame using a high-speed sCMOS detector and data were collected at 2Hz for 90-120 minutes.

##### Filleted animals

P2 stage pupae (Bainbridge and Bownes, 1981) were cold-anesthetized, washed in PBS, and dissected as for immunohistochemistry. The brain was removed by severing ventral nerve cord (VNC) projections and removing the brain and VNC. Once filleted, the PBS was replaced with 1mL of Schneider’s insect medium (SIM, Sigma Aldrich). Imaging was performed for 30 minutes on a Nikon SMZ25 stereomicroscope with a 1x/0.3 NA objective and sampled at 2Hz. After the first 5 minutes, 1 μM CCAP solution in SIM was added to the bath, for a final effective concentration of 0.5 μM, and image collection continued for the remaining 25 minutes. The vehicle control was SIM without CCAP. Ca^++^-free controls were conducted with SIM without Ca^++^ (Sigma Aldrich). CCAP (BACHEM, Torrence CA) stock solutions were prepared by dilution to 1mM in water.

##### Dual-channel imaging

For experiments requiring GCaMP6s and jRGECO1a fluorescence, animals were prepared as intact animals above and imaging was conducted on a Nikon Ti epifluorescence microscope with a 10X/0.5 NA air objective, Cairn twin-cam, two PCO edge 4.2 sCMOS cameras, and filters for GFP and RFP. Images were acquired at 2Hz for 60-90 minutes.

##### Image processing

Unless otherwise noted, all live experimental image series were background subtracted in Fiji ImageJ2 (Schindelin et al., 2012). Where values of length and fluorescence are indicated, the line tool was used to measure intensity and length for muscles 1, 2, 3, 5, 12, 13, 15, 22, 26, and 28 in hemisegments 3, 4, and 5 for each of the ecdysis sequence Phases at the onset of muscle activity (visible fluorescence), the peak of the muscle activity (brightest fluorescence), and the offset of muscle activity (fluorescence returns to baseline). Data were collected sub-saturation but most movies and figures presented are contrast-enhanced to aid the eye. Ca^++^ traces were extracted from ROIs defined as the entire animal excluding the puparium, or as single muscles/neurons where indicated, and normalized to F0, defined here as the average fluorescence intensity over the first 50 frames in the image series. Raster plots were generated from activity peaks identified from Ca^++^ traces using a peak finding algorithm in python with manually-determined thresholds.

#### Behavior annotation

A subset of 4 animals from the total *hlk*>GCaMP6s experimental dataset (N=16) were annotated frame-by-frame by expert raters to identify the muscles newly activated in each frame. These data were used to determine the frequency and order of activation for each muscle.

Movements were annotated manually for these 4 animals plus one more by visual inspection of muscle activity pattern and deflections of the body wall with respect to the puparium from raw image series (before background subtraction) that were contrast-enhanced to allow visualization of autofluorescence from the body wall. Bouts were identified manually from a subset of 10 *hlk*>GCaMP6s animals as the start of a sequence of muscle activations flanked on either side by ≥ 2 frames of no new activations, which were defined as inter-bout intervals.

### QUANTIFICATION AND STATISTICAL ANALYSIS

#### Sample calculations and general statistics

Sample sizes were predetermined using power calculations with 85% power from ecdysis phase 2 behavior duration data presented in Diao et al (2017). N calculated for suppression of the CCAP, non-ETHRA population vs WT is 6 animals and N calculated for suppressing CCAP/ETHRA population vs WT is 12 animals. Sample sizes in the current paper are listed with each figure and in Table S1 and range from 10 to 16 with the exception of manually annotated data, for which sample sizes were not statistically predetermined. Statistical analyses were performed using Graphpad Prism 8 or python 3.7 (https://anaconda.com, vers 2-2.4.0). Individual statistical methods and parameters are reported in the figure legends.

#### Directionality analysis

To find direction of activation (A->P or P->A), (ref. Fig 1K), we first computed a maximum intensity projection (MIP) along the x (horizontal) axis of the *hlk*>GCaMP6s image. For every frame, the MIP was a 1D vector of length H for an HxW image. Then we computed the location of the mode of the MIP vector on the y axis. An intensity mode indicates the location of the most dominant muscle activation. Therefore, the locations of the modes give an estimate of the direction of motion during the muscle activity period. Note that, we are interested in finding the location of maximum motion, which is efficiently captured by the mode of the intensities, not by the mean. Once the intensity modes were identified for every frame using MIPs, we computed the number of modes above and below a threshold, indicating P->A and A->P motion, respectively. The threshold is automatically computed as the median of all modes. We then calculated the ratio of the number of modes below the median line to the number of modes above the median line and used this ratio to determine if the direction of activation is posteriorly or anteriorly directed.

#### Automated movement analysis

We used a convolutional neural network (CNN) to automatically estimate the type of motion in each *hlk*>GCaMP6s image frame. The network model is a modified version of Inception-v3 (Szegedy, 2015). To fit the model in available GPU memory based on the image size, we removed the final Dense layer, and replaced it with a GlobalAveragePooling2D layer. In addition, a dropout layer was also added to introduce stochasticity into the model to avoid overfitting. There were total 21.8M parameters in the model. The detailed network is described in the train.py provided as supplementary material. The model was trained to predict 9 motions (Anterior Compression, Posterior Swing, Brace, Crunch, Rolling Contraction, Lift, Posterior Contraction, Rest, Swing) from the *hlk*>GCaMP6s images of 5 animals. For each animal, the images were resized to 403×129xN, where N was the variable number of frames based on the duration of the phases. Each frame was obtained at 2Hz. Every frame, starting from phase 1, was assigned to one of the above-mentioned motions, and was derived by consensus of 3 expert raters. Since the motion cannot possibly be derived by looking at a single frame, we used total 25 frames (12 previous and 12 following) to estimate the motion at one frame. Therefore, the training data for each frame consisted of a 403×129×25 matrix. The CNN model then trained by predicting the motion of the center (13^th^) frame.

Since there were only 5 animals available for training, we used all possible overlapping frames to augment the training data. Categorical cross-entropy with the Adam (Kingma, 2015) optimizer was used with learning rate of 0.0001 to obtain a probabilistic estimation of motion for each frame. An early termination criterion was used to prevent the model from overfitting, where the training was stopped if the categorical cross-entropy did not increase for 10 consecutive training epochs. Fig S3(A) shows the accuracy of motion prediction averaged over all available frames for each of the five training animals, in a leave-one-out cross validation. The final (magenta) curve shows the training accuracy where frames from all 5 animals were aggregated and then the model was trained on the aggregated frames. Similar accuracy between cross-validation and aggregated training shows that the CNN model did not overfit the data. However, to use information from all training animals, we applied the aggregated trained model (i.e., magenta) on the remaining 11 animals. For every frame of a test animal image, the motion with maximum probability was used as the final motion. For better accuracy, the predicted motions of the 11 animals were corrected by an expert rater.

#### Muscle activation order analysis

The annotated muscle datasets were sorted by frame number and then by muscle. Then, cells were concatenated into strings by bout, movement, or syllable, and assessed pairwise via the python SequenceMatcher algorithm to score for similarity in string order. The movements in each bout/phase were simplified to the most frequently occurring series in the real data (P1 bout: Lift, Rolling Contraction; P2 bout: Swing, then Swing, Brace; P3 bout: block 1 - Crunch, Brace, Anterior Compression, Posterior Gap, Posterior Swing, block 2 - Crunch, Brace, Anterior Compression, Posterior Gap, Brace). Documentation for this algorithm is provided by the difflib library (https://docs.python.org/3/library/difflib.html).

### DATA AND SOFTWARE AVAILABILITY

The data that support the findings of this study are available from the corresponding author upon reasonable request and code is posted to https://github.com/BenjaminHWhite.

## Supplemental Information

### Supplemental Tables

**Table S1. Pupal Neuromuscular Anatomy.**

**Table S2. Pupal Ecdysis Movement Components.**

**Table S3. Variability of Phase Parameters.**

### Multimedia Files

**Movie S1. Behavior at Single-Cell Resolution – Dorsal View.**

**Related to Figure 1.**

Dorsal view of a P1 bout with Lift and RollCon movements alternating bilaterally in a *hlk*>GCaMP6s animal; data were sampled at 2Hz and sped up to 10 fps. Scale bars, 250 μm. Time, seconds (s).

**Movie S2. Behavior at Single-Cell Resolution – Lateral View.**

**Related to Figure 1.**

Lateral view of a P1 bout with Lift and RollCon movements in a *hlk*>GCaMP6s animal; data were sampled at 2Hz and sped up to 10 fps. Scale bars, 250 μm. Time, seconds (s).

**Movie S3. Stochastic P0 Muscle Activity.**

**Related to Figure 2.**

Lateral view of P0 in a *hlk*>GCaMP6s animal; data were sampled at 2Hz and sped up to 20 fps. Scale bars, 250 μm. Time, seconds (s).

**Movie S4. The 8 Elementary Movements.**

**Related to Figure 3.**

Lateral view of each of the independent movements in a *hlk*>GCaMP6s animal; data were sampled at 2Hz and sped up to 10 fps, except the posterior contraction, which is 5 fps. Scale bars, 250 μm. Time, seconds (s).

**Movie S5. Attempted Swing of N_CCAP_-suppressed Pupa.**

**Related to Figure 6.**

Lateral view of N_CCAP_ bouts including attempted swing movement (top), with wildtype P2 bouts for comparison (bottom); data were sampled at 2Hz and sped up to 5 fps. Scale bars, 250 μm. Time, seconds (s).

**Movie S6. CCAP Activation of M21-23.**

**Related to Figure 6.**

Live fillet of a *hlk*>GCaMP6s animal before and after bath application of synthetic CCAP; data were sampled at 2Hz and sped up to 10 fps. Scale bars, 250 μm. Time, seconds (s).

