## Supplemental Figures and Tables for "Behavior emerges from unstructured muscle activity in response to neuromodulation"

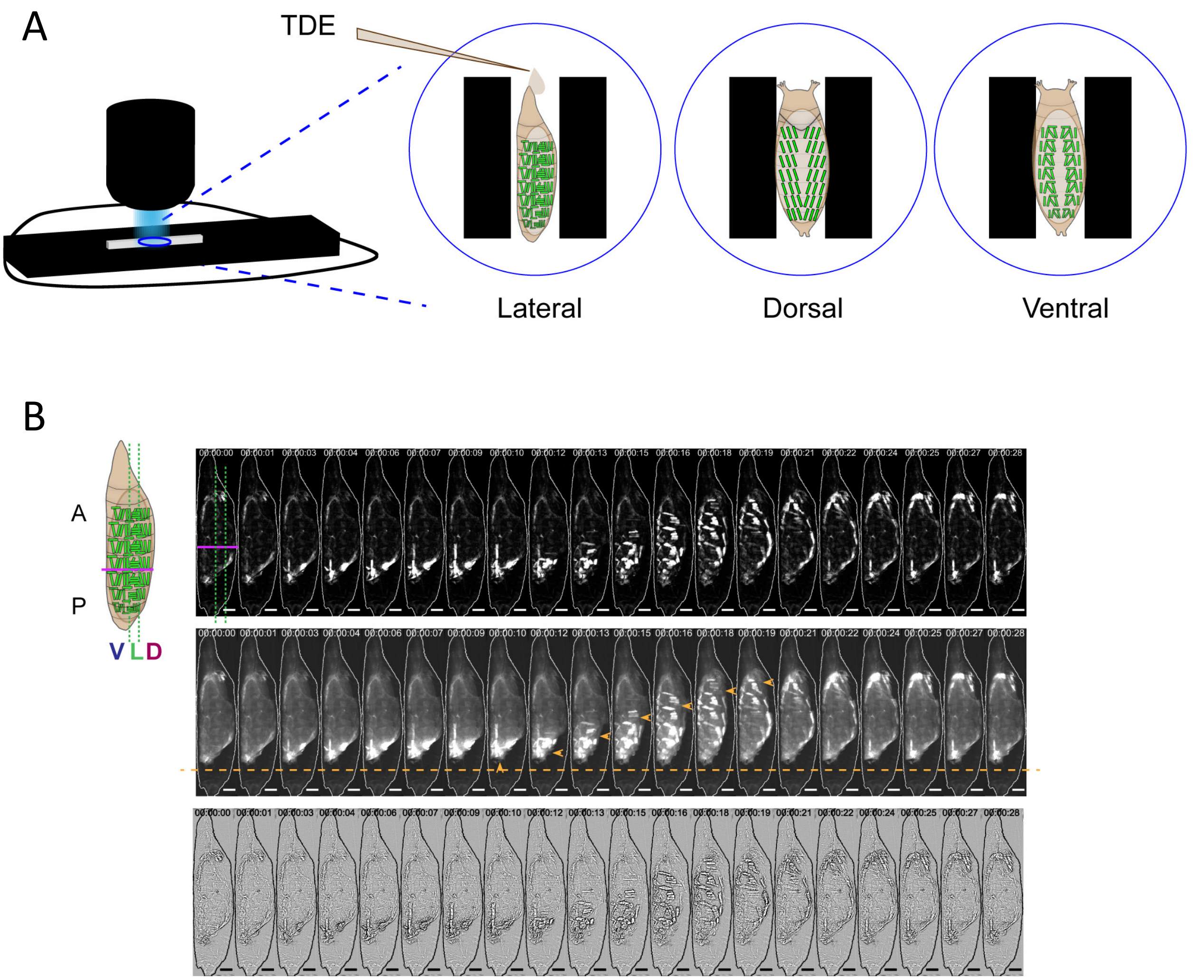


**Figure S1. Paradigm for Imaging Muscle Activity and Body Wall Movements. Related to Figure 1.**

**(A)** Schematic of imaging workflow for single-muscle resolution data collection. Epi-illuminated stereomicroscope on the left was used to image coverslips containing TDE-filled chambers and animals oriented as shown enlarged on the right for live imaging.

**(B)** Image analysis for muscle activity and body wall movements. On left, schematic of *hlk*>GCaMP6s pupa (muscles in green) from the lateral view. V and D: ventral and dorsal compartments, respectively, with lateral region (L) flanked by green dotted lines. Magenta, A-P boundary. On right, three representations of an image montage from a P1 activity bout taken from the lateral view: (top) background-subtracted and scaled to show muscle activation; (middle) not background-subtracted to show abdominal body wall deflections (orange arrowheads) via autofluorescence; and (bottom) Laplace transformed to enhance body wall deflections. Orange line, posterior limit of body. Scale bars, 250 µm.

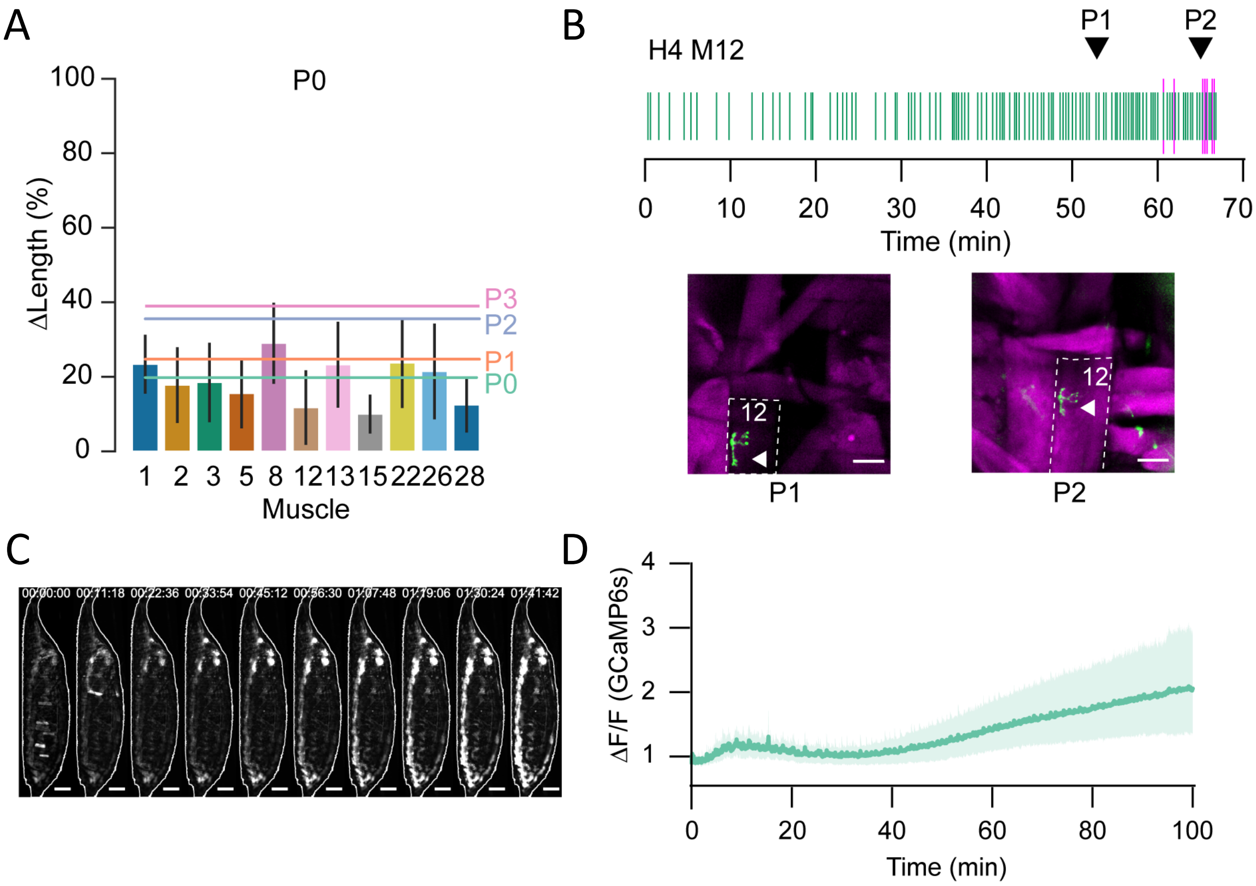


**Figure S2. Simultaneous Imaging of Muscle Activity with Synaptic or Proprioceptive Activity. Related to Figure 2.**

**(A)** Bar plots with percent change in length (ΔL/L) plotted as mean ± SD for muscles measured during P0. Colored lines represent mean length changes of all muscles for each phase (green, P0; orange, P1; blue, P2; pink, P3). N=11 muscles, 3 hemisegments, 8 animals.

**(B)** Raster plot of peaks in muscle M12 $Ca^{++}$ activity (magenta) measured with *hlk*>jRGECO and M12 NMJ activity (green) measured with Syt-GCaMP6s during P0, P1, and P2, as indicated. Coincidence in NMJ and muscle activity begins near the onset of P2. Images of M12 (outlined) in hemisegment 4 showing that during P1 (left), M12 muscle activity (magenta) is low in response to NMJ activation (arrowhead, green), whereas in P2 (right) activity is high. Scale bars, 50 µm.

**(C)** Image montage of bulk muscle activity in a *hlk*>GCaMP6s animal with class I da proprioceptors suppressed by two copies of Kir2.1. Scale bars, 250 µm.

**(D)** Bulk $Ca^{++}$ fluorescence trace (mean ± SD) for N=10 animals like the one shown in (C). Stochastic activity is lost, followed by death before ecdysis onset.

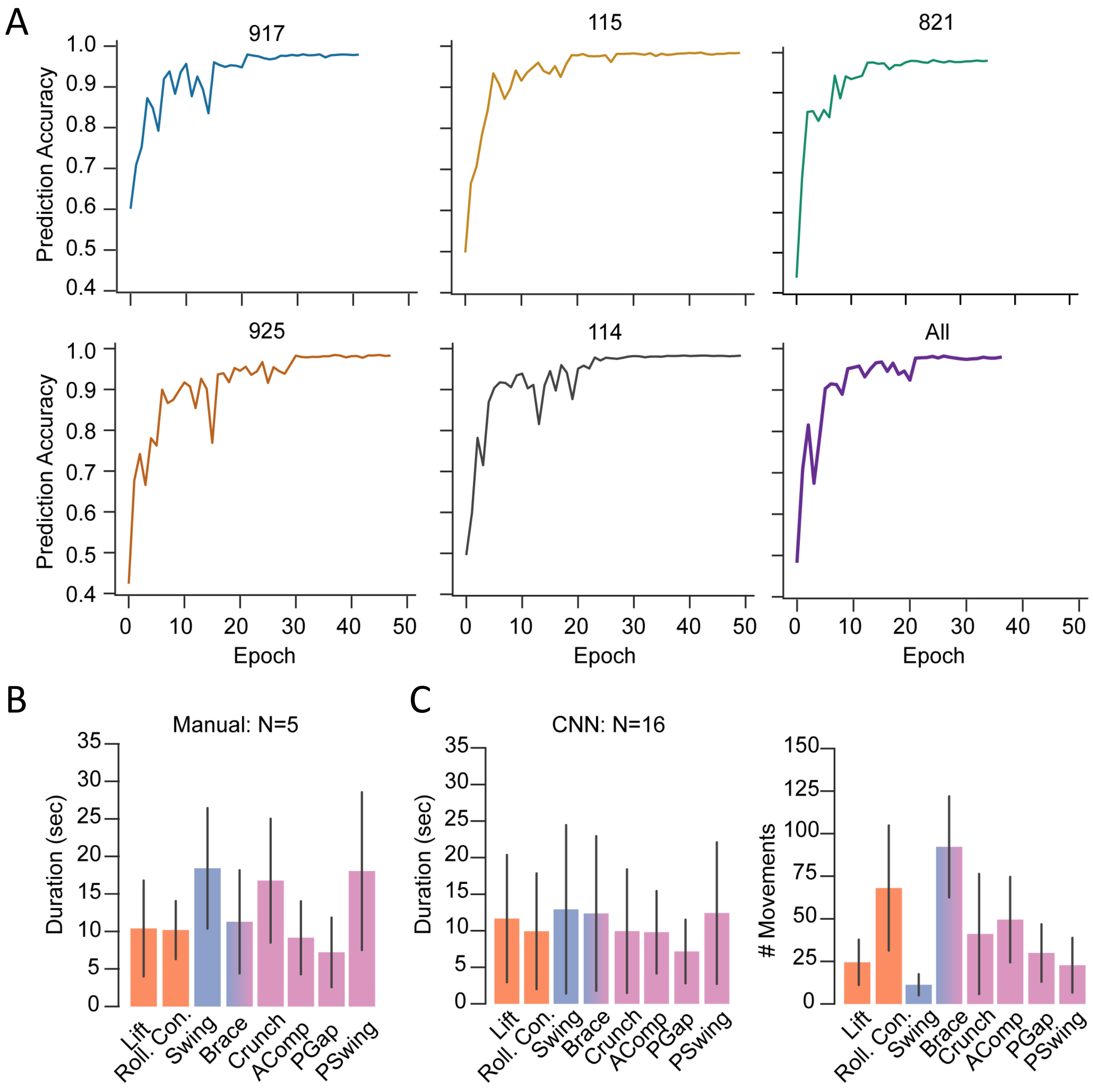


**Figure S3. Analysis of Movement Using a Convolutional Neural Network.**
**Related to Figure 3.**
**(A)** Graphs showing the CNN's accuracy of movement prediction per epoch for the five training samples (917, 115, 821, 925, 114) from a "leave-one-out" cross validation. The last graph (labeled "All") shows the training accuracy using all 5 training samples, which represents the model used on the other animals (N=16).
**(B)** Movement durations shown as mean ± SD for the five manually annotated animals.
**(C)** Movement durations (left) and numbers (right) as determined by the CNN (mean ± SD). N=16 animals.

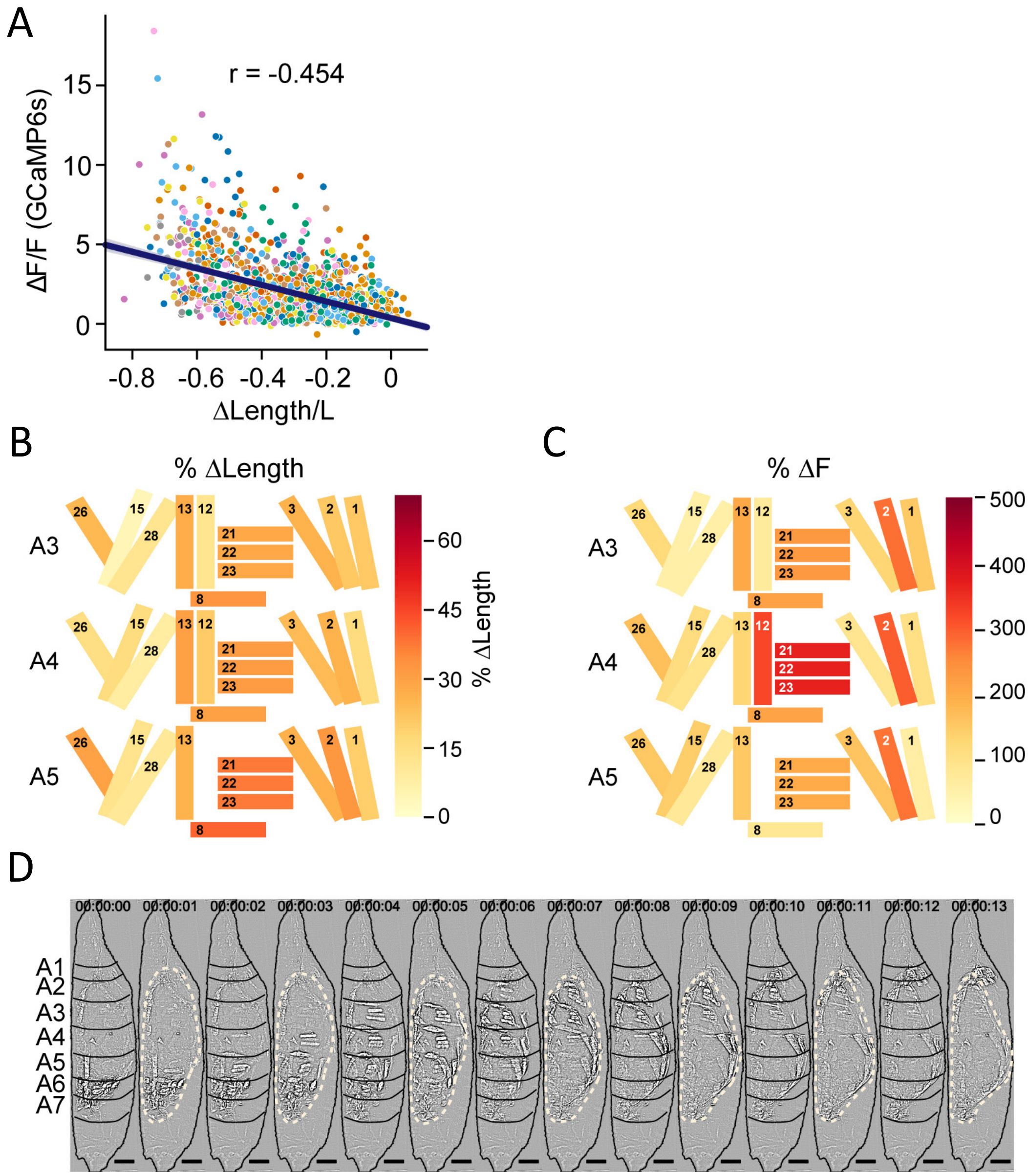


**Figure S4. Muscle Length and Fluorescence Changes Underlying P1 Movements. Related to Figure 5.**

**(A)** Scatter plot of normalized change in length (ΔL/L) vs normalized change in fluorescence (ΔF/F) for muscles from eight animals (colored by animal). Line shows Pearson correlation; r indicated.

**(B-C)** Heatmaps of changes in muscle properties during contraction of the indicated muscles in hemisegments A3-A5 during P1 RollCon activity. (**B**) % change in muscle fiber length (ΔL/L). **(C)** % change in GCaMP6s fluorescence (ΔF/F). In both cases, changes were measured at the onset of activity and at maximum activity ($(t_1-t_0)/t_0$) for N=24 animals.

**(D)** Image montage of muscle activity comprising a P1 RollCon after Laplace transformation to show body wall distortion during movement (black lines, segments; beige dotted lines, pupal body). Scale bars, 250 µm.

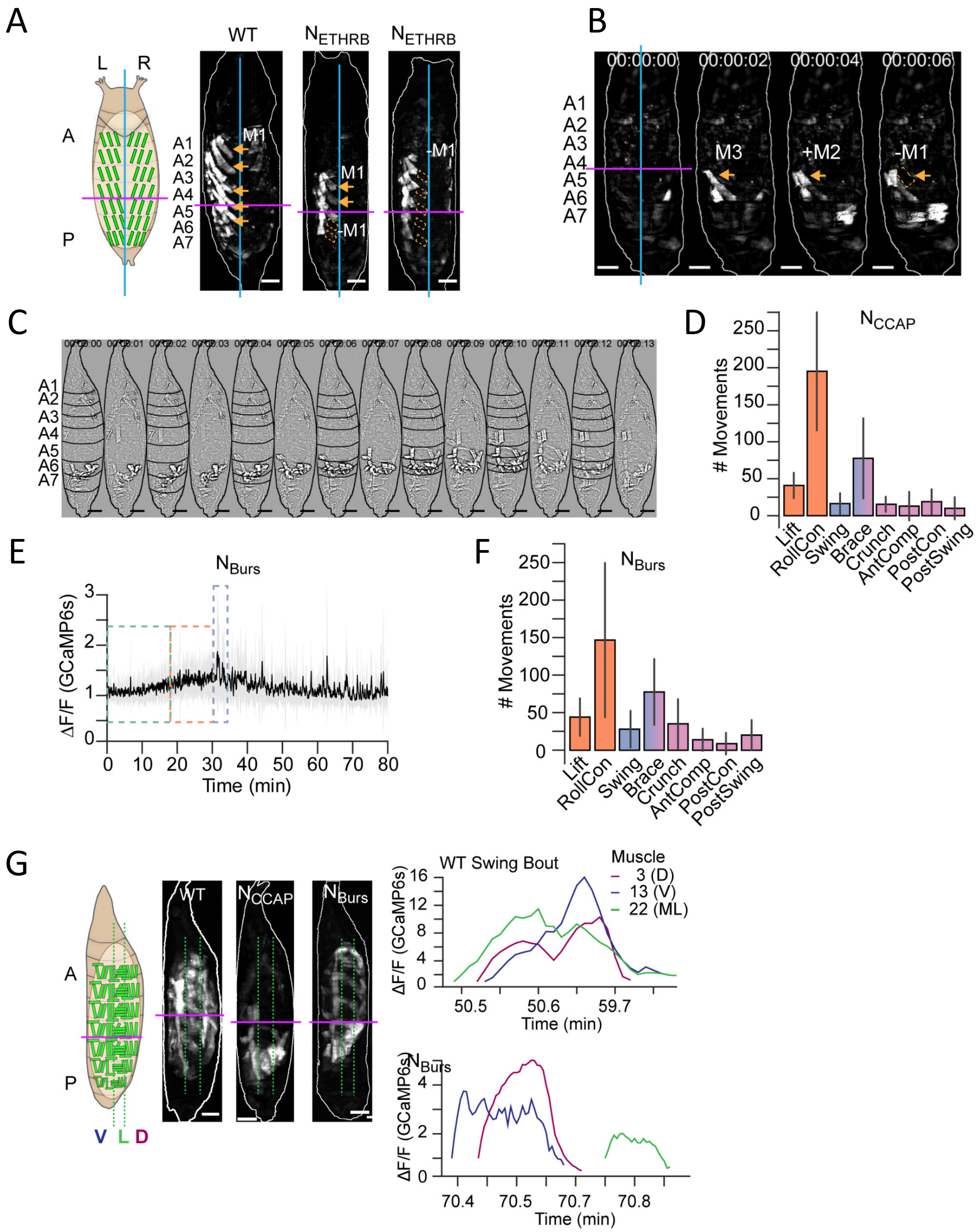


**Figure S5. Neuromodulator Action on Syllables and Movements.**
**Related to Figure 6.**
**(A)** Schematic on left shows L-R and A-P axes with lines indicating the dorsal midline (cyan), and the A-P boundary between segments A4 and A5 (magenta). Dorsal images of a left Swing

from wildtype (left) and two $N_{ETHRB}$-suppressed animals (middle and right) showing disrupted M1 activity. Orange arrows, M1 activity; dotted boxes, missing M1 activity. Scale bars, 250 µm.

**(B)** Image montage (dorsal view) of muscle activity comprising the single, partial "P2" swing of an animal with suppressed $N_{CCAP}$. PME4 incompletely forms (orange arrowheads) and the P-to-A activity reaches only segment 4. Left image indicates the dorsal midline (green line) and P-A boundary (magenta line). Scale bars, 250 µm.

**(C)** Image montage (lateral view) of muscle activity comprising the single, swing-like movement of an animal with suppressed $N_{CCAP}$ after Laplace transformation to show segmental compressions. Black lines, segmental boundaries. Scale bars, 250 µm.

**(D)** The number of movements in $N_{CCAP}$-suppressed animals as detected by CNN and plotted as mean (± SD). N=10 animals.

**(E)** Bulk $Ca^{++}$ activity trace (lateral view) for an animal in which the Bursicon-expressing neurons ($N_{Burs}$) are suppressed. Mean activity (black) ± SD (gray) is shown for 10 animals. Dotted outlines indicate identifiable phases, with P2 consisting of a single attempted swing, though subsequent attempts and some P3-like activity follows.

**(F)** The number of movements as detected by CNN in $N_{Burs}$ animals plotted as mean (±) SD. N=10 animals.

**(G)** Images on the left show muscle activity comprising Swing, or swing-like, movements in wildtype (WT), $N_{CCAP}$-suppressed, and $N_{Burs}$-suppressed animals (schematic as in Fig. 1J); $Ca^{++}$ traces on right show activity of muscles M3, M13, and M22 in HS A4 from a single Swing bout in a wildtype animal (top) compared with traces for the same muscles from the swing-like bout in a $N_{Burs}$-suppressed animal. As for $N_{CCAP}$-suppressed animals, co-incidence across the dorsal (D), lateral (L), and ventral (V) compartments is lost in $N_{Burs}$-suppressed animals. Scale bars, 250 µm.

**Table S1. Pupal Neuromuscular Anatomy.**

| Spatial Group (N=17) | Muscle/ MN-IB (N=17) | CCAP-R+ MN (N=11) | Nerve | Segments (N=17) | CCAP-R+ Muscle (N=5) |
|---|---|---|---|---|---|
| D/DL | 1 | + | $ISN^{DM}$ | A1-A7 | - |
| D/DL | 2 | + | $ISN^{DM}$ | A1-A7 | - |
| D/DL | 3 | + | $ISN^{DM}$ | A1-A7 | - |
| D/DL | 9 | + | $ISN^{DM}$ | A1-A7 | - |
| D/DL | 10 | + | $ISN^{DM}$ | A1-A7 | - |
| D/DL | 4* | - | $ISN^{DM}$ | A1-A3 | - |
| V/VL | 12* | - | ISNb | A1-A4 | - |
| V/VL | 13 | + | ISNb | A1-A7 | - |
| V/VO | 28 | + | ISNb | A1-A7 | - |
| V/VO | 15 | + | ISNd | A1-A7 | - |
| L/DO | 5* | - | SNa | A1-A4 | - |
| L/TR | 8 | - | SNa | A1-A7 | - |
| L/TR | 21/22 | - | SNa | A1-A7 | + |
| L/TR | 22/23 | - | SNa | A1-A7 | + |
| L/TR | 23/24 | - | SNa | A1-A7 | + |
| V/TR | 25 | - | TN | A1-A7 | - |
| V/VA | 26 | - | SNc | A1-A7 | - |

a. Abbreviations: D, dorsal; L, lateral; V, ventral; DL, dorsal longitudinal; VL, ventral longitudinal; VO, ventral oblique; DO, dorsal oblique; TR, transverse; VA, ventral acute

b. * Degrades prior to pupal ecdysis in segments posterior to HS3 or HS4

c. N is the number of animals

**Table S2. Pupal Ecdysis Movement Components.**

| Movements | - | Lift | RollCon | Swing | Brace | Crunch | AntComp | PostCon | PostSwing |
|---|---|---|---|---|---|---|---|---|---|
| Compartments | - | P D/L/V | A/P D/L/V | A/P D/L/V | A/P L | A/P D/L/V | A D/L/V | P D/L/V | P D/L/V |
| Syllables | PME 2, 3, 5, 6 All M | PME 1, 2, 3, 4, 6 M8, M15 | PME 2, 6 M2, M8 | PME 1, 2, 3, 4, 6 M15, M12 | PME 2 M8 | PME 1, 2, 6 M2, M8, M13 | PME 3, 5, 6, 7, 8 M1, M12 | PME 2, 3, 6 M8, M26 | PME 1, 3, 6 M15 |
| L-R Rhythm | L-R | - | L-R | L-R | L-R | L-R | - | L-R | L-R |
| A-P Rhythm | - | P-A | P-A | P-A | A-P | A-P P-A | A-P | - | A-P |
| Phase | P0 | P1 | | P2 | P2/P3 | P3 | | | |
| Function | - | Fragment Trachea | | Evert Head, Shed Trachea | | Elongate Appendages | | | |

a. Abbreviations: D, dorsal; L, lateral; V, ventral; A, anterior; P, posterior; L-R, left-right; A-P, anterior-posterior; PME, pupal muscle ensemble

b. Individual muscles indicated as ‘M’ followed by the number

**Table S3. Variability of Phase Parameters.**

| Phase | Mean ± SD | Mean CV ± SD | N |
|---|---|---|---|
| **P0** | | | |
| Phase Duration (min) | - | - | 16 |
| Bout # | - | - | 10 |
| Bout Duration (sec) | 34.42 ± 9.45 | 0.41 ± 0.15 | 10 |
| IBI Duration (sec) | 8.25 ± 3.33 | 0.82 ± 0.33 | 10 |
| **P1** | | | |
| Phase Duration (min) | 12.63 ± 3.45 | 0.27 | 16 |
| Bout # | 25.1 ± 5.17 | 0.51 ± 0.093 | 10 |
| Bout Duration (sec) | 31.43 ± 4.63 | 0.36 ± 0.061 | 10 |
| IBI Duration (sec) | 3.43 ± 1.32 | 0.72 ± 0.14 | 10 |
| **P2** | | | |
| Phase Duration (min) | 5.68 ± 1.02 | 0.18 | 16 |
| Bout # | 9.1 ± 0.88 | 0.51 ± 0.099 | 10 |
| Bout Duration (sec) | 31.23 ± 4.01 | 0.32 ± 0.066 | 10 |
| IBI Duration (sec) | 5.33 ± 1.67 | 0.63 ± 0.12 | 10 |
| **P3** | | | |
| Phase Duration (min) | - | - | 16 |
| Bout # | - | - | 10 |
| Bout Duration (sec) | 85.21 ± 16.32 | 0.38 ± 0.058 | 10 |
| IBI Duration (sec) | 6.89 ± 1.87 | 0.62 ± 0.1 | 10 |

a. Abbreviations: SD, standard deviation; CV, coefficient of variation

b. N is the number of animals
